# The methionine cycle sustains the mesenchymal phenotype of tumor cells and drives pancreatic cancer progression

**DOI:** 10.64898/2026.11.05.764593

**Authors:** E. Jaune-Pons, G. Efthymiou, E. Metay, V. Gouirand, Q. Da-Costa, P. Finetti, S-S. Tubiana, C. Neuzillet, E. Lohmann, S. Djefaflia-Beloribi, E. Mamessier, P. Grenot, N. Dusetti, R. Tomasini, F. Bertucci, F. Guillaumond, S Vasseur

## Abstract

Mesenchymal traits are associated with poor prognosis in Pancreatic Ductal Adenocarcinoma (PDA), but direct targeting of mesenchymal tumor cells remains a challenge. Metabolic profiling of tumor cells isolated from murine PDA and liver metastases (LM) revealed that LM rely more on the methionine cycle than does primary PDA. This metabolic shift was specifically associated with increased activation of the methionine cycle in mesenchymal tumor cells compared to differentiated tumor cells. We show that the targeting of the key enzyme of the methionine cycle S-adenosylhomocysteine hydrolase (SAHH) in mesenchymal cells lowers the levels of homocysteine (Hcy) and glutathione, resulting in reduced antioxidant capacity, deregulated intracellular calcium homeostasis, and ultimately loss of mesenchymal markers. Together, these events decrease cell proliferation and invasion, blunting the tumorigenic and metastatic potential of SAHH-deficient cells, eventually impeding tumor growth and progression of the metastatic disease. Clinically, the high expression of SAHH in patients with mesenchymal PDA is correlated with poor survival, positioning SAHH as a potential therapeutic target to impede the pro-metastatic, mesenchymal phenotype of PDA.

## INTRODUCTION

Pancreatic ductal adenocarcinoma (PDA) is globally increasing, presents the poorest prognosis among all gastrointestinal cancers, and is the seventh most common cause of cancer-related deaths^1^. Most patients are diagnosed when the disease has progressed to a metastatic stage when the tumor is unresectable and poorly responsive to conventional therapies. Pancreatic tumor cells often disseminate to distant organs such as the liver (75-95%), the peritoneum (40-55%), the abdominal lymph nodes (40%), and the lungs (45-50%) with a seeding occurring before or concomitantly to tumor formation at the primary site^2^. In aggressive PDA, tumor cells are embedded in a dense stroma which limits their access to oxygen and nutrients from the bloodstream and forces them to adapt their metabolism to face these stringent microenvironmental conditions. Once PDA has reached an advanced stage, malignant cells from the primary tumor display metabolic traits that might contribute to metastatic spread. Upon metastatic progression, cancer cells gradually lose their epithelial features and gain mesenchymal characteristics via epithelial to mesenchymal transition (EMT)^3–5^. Indeed, these cells progress in a stepwise manner through EMT by co-expressing epithelial and mesenchymal markers at various degrees corresponding to partial/hybrid (h), or complete (c) EMT states that correlate to a range of invasive and migratory tumor cell behaviors^6–9^. Interestingly, patients with a predominantly epithelial PDA exhibit a better overall survival^10^ suggesting that abrogation of mesenchymal features in tumor cells may improve patient outcomes.

Throughout cancer progression, tumor cells acquire distinct metabolic features that promote the mesenchymal phenotype and drive their metastatic potential. For example, high levels of D-2HG (D-2 Hydroxyglutarate) in colorectal cancer cells are associated with increased expression of *ZEB1*, a master inducer of EMT^11^, while the oxidative pentose phosphate pathway via the enzyme 6-phosphogluconate dehydrogenase (PGD) promotes the expansion of metastatic subclones in PDA via epigenetic alterations^12^. This was supported by the unexpected finding that glutamine deprivation depriving of PDA cells resulted in SLUG-mediated EMT induction via MEK/ERK signaling and subsequent promotion of metastasis^4^. Interestingly, dependency on glutaminase 1 (GLS1) for redox balance via glutathione synthesis was associated with mesenchymal breast and lung tumors^13^. Conversely, the EMT transcription factors TWIST and SNAI2 downregulate the succinate dehydrogenase B and C thereby reducing mitochondrial mass and respiration^14^, while increased glycolysis-derived lactate was observed in the PDA cell line Panc-1 when subjected to TNFα-or TGFβ-induced EMT^15^. From the above it is clear that the elucidation of the mechanisms by which metabolic rewiring promotes mesenchymal features may provide attractive approaches to impede PDA progression and subsequent metastatic dissemination.

Here, we present a matched-pair comparison of the metabolic transcriptome of PDA and liver metastasis (LM). This analysis shows an upregulation of transcripts coding for enzymes of the methionine cycle in LM, which correlates with the acquisition of a mesenchymal phenotype. We also demonstrate that mesenchymal PDA cells with an activated methionine cycle display increased ability to balance high ROS levels. Within this setting, we discovered that the enzyme S-adenosylhomocysteine hydrolase (SAHH), that catalyzes the reversible hydrolysis of S-adenosylhomocysteine (SAH) to adenosine and homocysteine (Hcy), is essential for mesenchymal PDA cells to maintain redox potential and calcium homeostasis and sustain their proliferative and invasive capacities. Indeed, the targeting of SAHH drastically attenuates the mesenchymal features of PDA cells, resulting in restricted tumor growth and diminished capacity of LM formation for tumors with a strong mesenchymal phenotype.

## RESULTS

### Mesenchymal tumor cells display high expression of methionine cycle enzymes

To explore the metabolic preferences of PDA tumor cells at the primary site and after metastatic dissemination into the liver, we compared the transcriptomic profiles of micro-dissected tumor cell areas from spontaneous PDA and LM collected from *LSL-Kras^G12D/+^; Ink4a/Arf^fl/fl^; Pdx1-Cre* (KIC) mice^16^ (**Supp Figure 1A**). We previously showed that PDA tumor cells rely on specific amino-acid pathways when they are in a stressful environment or when they acquire specific oncogenic functions. Indeed, under nutrient stress they rely on macropinocytosis to take up collagen from which they use proline for metabolic purposes^17^, and they activate branched-chain amino acid and ketone body metabolisms to evade the primary tumor and form liver metastasis^18^. Therefore, we performed Significance Analysis of Microarrays on our dataset, focusing on genes involved in amino acid metabolism (**Supp Figure 1B, Supp Table 1**). We identified 81 significantly upregulated genes in LM compared to PDA (**Supp Figure 1B, Supp Table 2**). These genes clustered in 11 metabolic pathways, of which the methionine, cysteine and glutathione metabolic pathways were the most significantly enriched in LM compared to PDA (**Figure 1A, Supp Table 3**). The genes coding for enzymes of the methionine cycle (*e.g.*, *Mat1a,* methionine adenosyl transferase1a; *Ahcy,* adenosyl homocysteinase; *Gnmt,* glycine N-methyltransferase; *Bhmt2,* betaine-homocysteine *S*-methyltransferase2), of the transsulfuration pathway which allows the conversion of Hcy to cysteine (*e.g.*, *Cbs,* cystathionine β-synthase; *Cth*, cystathionase), and of glutathione (GSH) synthesis (*e.g.*, *Gclc,* glutamate-cysteine ligase catalytic subunit) were highly upregulated in LM compared to PDA (**Figures 1B, C, Supp Figure 1C**). The protein levels of SAHH (coded by *Ahcy*), and CTH were increased in LM tumor cells compared to the PDA tumor cells, while MAT1A and DNMT1 were uniformly abundant at both sites (**Figure 1D**). Interestingly, both SAHH and CTH, that are the intersection of the methionine cycle and cysteine biosynthesis (**Supp Figure 1C**), localized in disseminated tumor cells in PDA and in LM rather than in well differentiated PDA cells within tumor glands (**Figure 1E**).

**Figure 1.**
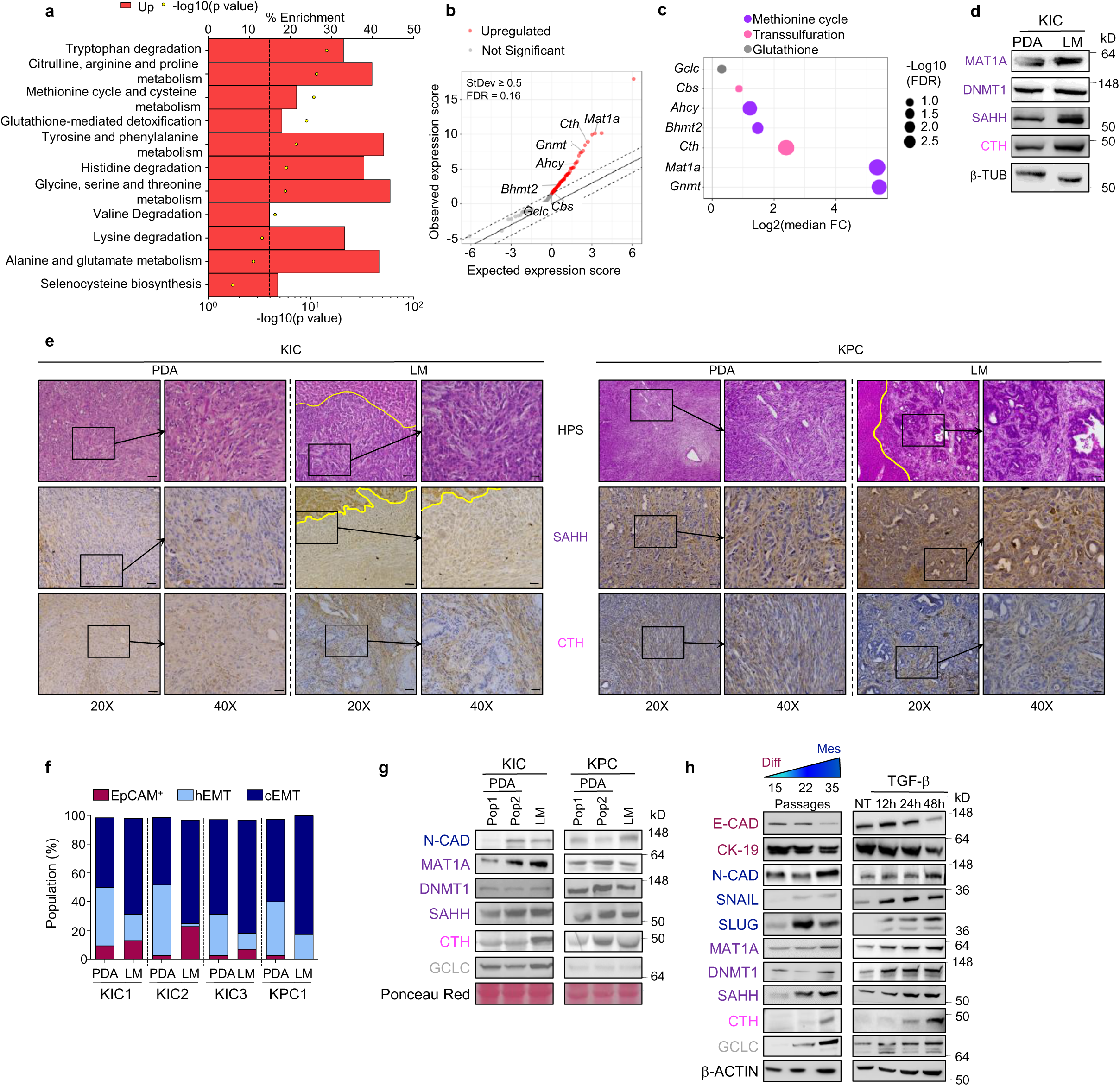
Methionine cycle enzymes are increased in liver metastasis compared to PDA and correlated with the expression of EMT markers. **a**, Enrichment analysis based on 81 genes upregulated genes in LM relative to PDA. The dots represent the enrichment p-value (x-axis) of each pathway (n=4 mice). The black dotted line indicates p-value=0.0001. b, Expression of the 81 genes in LM relative to PDA based on significant analysis of microarray (SAM). The indicated genes are involved in the methionine cycle, transsulfuration and glutathione pathways. c, Dot plot of the seven genes (from b) involved in the methionine cycle (purple), transsulfuration (pink) and glutathione (grey) pathways. Dot size is proportional to -log10(FDR). d, Comparison of the expression of enzymes between PDA and LM from KIC mice. Methionine cycle, purple; transsulfuration, pink; β-tubulin (β-TUB) was used as loading control. Representative immunoblots from n=2 independent experiments). e, Representative IHC images (HPS staining, SAHH and CTH immunostaining) from PDA and LM from KIC (left, n=2 mice) and KPC (right, n=2 mice). Yellow lines delineate LM within the liver tissue. Scale bars: 20X, 50 µm; 40X, 20 μm. f, Distribution of tumor cell populations from PDA and from the matched LM (n=3 KIC and n=1 KPC mice) into epithelial (EpCAM^+^), hybrid EMT (hEMT) and complete EMT (cEMT), based on flow cytometry analysis of epithelial and mesenchymal markers. g, WB on Pop1 and Pop2 tumor cells from the PDA and on tumor cells from the matched LM of KIC (left, n=2) and KPC (right, n=2) mice. EMT marker (N-CAD, blue); enzymes of the methionine cycle (purple), of the transsulfuration (pink), glutathione (grey) pathways. Representative immunoblots from n=3 independent experiments. Ponceau red was used as loading control. h, WB showing the expression of epithelial (red), EMT (blue) markers, of enzymes of the methionine cycle (purple), of the transsulfuration (pink), glutathione (grey) pathways in PK4A cells at early (Diff), intermediate, and late (Mes) passages (left), or after treatment with TGF-β (right). β-Actin was used as loading control.

Because EMT is typically associated with PDA metastatic progression^19,20^, we set out to determine the percentage of tumor cells at various EMT states in PDA and LM in the KIC and the *Pdx-1-Cre;LSL-Kras^G12D^;LSL-Trp53^R^*^172^*^H^* (KPC)^21^ mouse models of metastatic PDA. We screened these cells for expression of EMT surface markers such as the epithelial marker EpCAM (a cell adhesion molecule)^22^, CD106 (also known as VCAM1)^23^ associated with metastatic progression in gastric and breast cancers^24,25^, and CD51 (also known as integrin alpha V, ITGAV) that can propagate TGF-β-induced signaling to promote EMT^26,27^ (**Supp Figure 1D**). In both mouse models, we found that only a small fraction of PDA cells was EpCAM^+^ and that PDA displays a higher percentage of cells in a hEMT state (EpCAM^-^/CD51^-^/CD106^+/-^) than LM (**Figure 1F**). Interestingly, up to 69% of PDA cells were engaged in cEMT (EpCAM^-^/CD106^+/-^/CD51^+^), while this cEMT cell subset increased up to 80% in LM at the expense of the hEMT cell subset (**Figure 1F**). Of note, in KIC-derived LM, EpCAM^+^ cells were more abundant than in PDA, suggesting that a Mesenchymal-to-Epithelial Transition (MET) occurs in these metastatic foci. Immunohistochemistry (IHC) staining confirmed the presence of EpCAM^+^ areas as well as of mesenchymal CD51^+^ and N-cadherin^+^ (N-CAD^+^) cells in both primary and secondary tumor sites (**Supp Figure 1E**). We next investigated the levels of enzymes involved in the methionine cycle and the transsulfuration pathway in epithelial *versus (vs)* cEMT tumor cells using fluorescence-activated cell sorting (FACS). We sorted EpCAM^+^ and cEMT tumor cell subsets (**Supp Figure 1F**) and observed that EpCAM^+^ cells lost almost completely the epithelial marker after only 2 to 3 passages and shifted towards a mixed population 1 (Pop1) composed mostly of hEMT and to a lesser extent of cEMT cells (**Supp Figure 1G**). In addition, the cEMT cell subset evolved towards a population 2 (Pop2) composed of hEMT cells (around 20%), of EpCAM-re-expressing cells (in the KPC mouse model) and of up to 85% of cEMT cells (**Supp Figure 1G**). These results underline the transient nature of EpCAM and demonstrate the ability of tumor cells to switch from an epithelial to a mesenchymal state, and *vice versa*, rather than keeping a stable EMT phenotype. Interestingly, increased N-CAD levels in Pop2 and in LM tumor cells from KIC mice were correlated with higher levels of the methionine cycle enzymes MAT1A and SAHH, as well as of the transsulfuration and GSH metabolism enzymes CTH and GCLC, respectively (**Figure 1G)** as compared to Pop1 cells. In Pop1, Pop2 and LM cells from KPC mice, elevated N-CAD also correlated with high levels of MAT1A, DNMT, SAHH. While CTH was abundant in Pop2 and LM from KPC mice, no correlation was found between N-CAD and GCLC in these mice. To further explore the correlation between the EMT state of tumor cells and the levels of enzymes involved in the methionine cycle, transsulfuration and GSH metabolism, we isolated fresh PDA cells (namely PK4A cells) and cultured them over ten to twenty passages (**Supp Figure 1H left)**. The PK4A cells progressed from a differentiated (Diff) phenotype, characterized by at least 50% of cells in epithelial and hEMT states and by high levels of E-cadherin and CK19, to a mesenchymal (Mes) phenotype, mainly characterized by a cEMT state and by increased levels of N-CAD, and of the EMT-transcription factors SNAIL and SLUG^28–30^ (**Figure 1H left, Supp Figure 1I top**). We observed a progressive increase of MAT1A, DNMT1, SAHH as well as of CTH and GCLC while PDA cells progressed towards a cEMT state and expressed mesenchymal markers (**Figure 1H left**). Similar data were obtained using a model of TGFβ-induced EMT in PK4A cells at low passage (**Supp Figure 1H right, Figure 1H right, Supp Figure 1I bottom, and Supp Figure 1J**)^31^. Collectively, these findings highlight that PDA cells whether harboring an *in situ*-acquired EMT phenotype or submitted to a cell autonomous or a TGFβ-induced EMT program, upregulate their levels of enzymes of the methionine cycle, transsulfuration and GSH pathway.

### Mesenchymal PDA cell aggressiveness is associated with activation of the methionine cycle

Depending on the EMT state and the type of cancer, tumor cells exhibit distinct behaviors during cancer progression and metastasis. We therefore defined the respective tumorigenic and metastatic potential of PDA cells according to their EMT state. We showed that following an orthotopic injection of PDA cells in a syngeneic mouse model, tumors derived from Mes PK4A cells had an increased weight compared to those derived from Diff PK4A cells (**Figure 2A, Supp Figure 2A**). Interestingly, in Mes tumors, SAHH co-localized with N-cadherin and CD51 (**Supp Figure 2B**).

**Figure 2.**
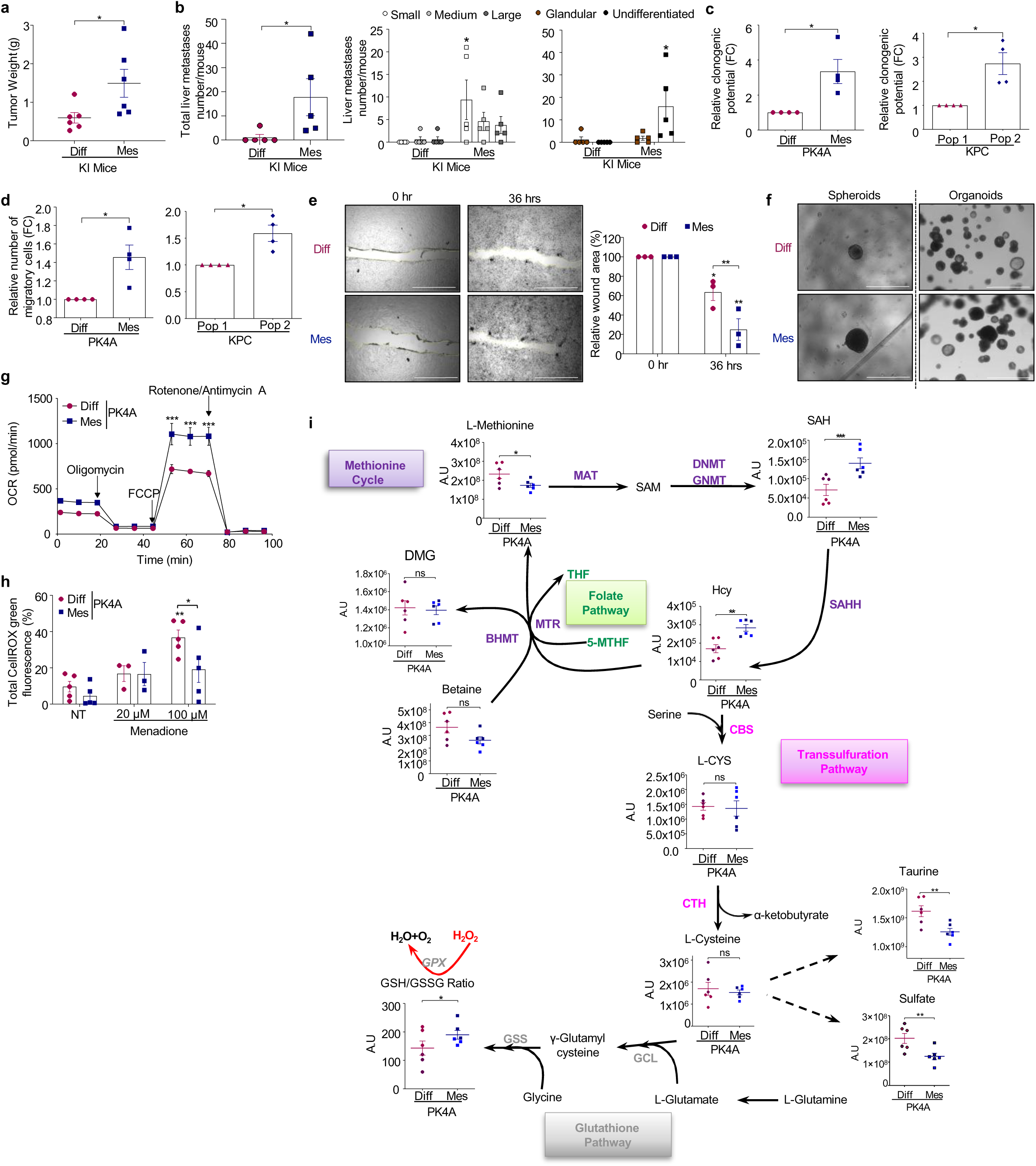
Mesenchymal PDA cells activate the methionine cycle and show aggressive oncogenic features and enhanced metabolic potential. **a,** Mean±SEM of the PDA weight in mice engrafted with Diff or Mes PK4A cells (n=6 mice/group). **b,** Metastasis formation after the implantation of Diff or Mes PK4A cells in the liver (n=5 mice/ group) and presented as the total number of metastases (left) or classified by size (center; white, small; grey, medium; black, large) and by their histological status (right; brown, glandular; black, undifferentiated). Data expressed as mean±SEM. **c,** Clonogenic potential of Diff and Mes PK4A cells (left) and KPC Pop1/2 cells (right) after 72 hrs, expressed as mean±SEM of the fold-change (FC) of Mes and Pop2 relative to Diff or Pop1 cells respectively from n=4 independent experiments. **d,** Migration of Diff and Mes PK4A cells (left) and KPC Pop1 and Pop2 cells (right) after 72hrs expressed as mean±SEM of the FC of Mes and Pop2 relative to Diff or Pop1 cells respectively from n=4 independent experiments. **e,** Representative images (left) and quantification (right) of the wound healing capacity of Diff and Mes PK4A cells after 36h. presented as mean±SEM of the percent reduction of the wound area at 0h from n=3 independent experiments. Scale bar, 2 mm. **f,** Representative images of Diff and Mes PK4A spheroids (left) and organoids (right) from n=3 independent experiments, spheroid assay; n=3 biological replicates, organoid assay. Scale bars, 1 mm. **g,** Oxygen consumption rate (OCR) in Diff and Mes PK4A cells expressed as mean±SEM from 3 independent experiments. **h,** Total ROS production by non-treated and menadione-treated Diff and Mes PK4A cells expressed as mean±SEM percentage of fluorescence from n≥3 independent experiments. **i,** Levels of metabolic intermediates of the methionine cycle, transsulfuration and glutathione pathways in Diff and Mes PK4A cells expressed as mean±SEM in Arbitrary Units (A.U) from 2 independent experiments performed in triplicates. Methionine adenosyltransferase (MAT); S-adenosyl methionine (SAM); DNA methyltransferase (DNMT); Glycine N-methyltransferase (GNMT); S-adenosyl homocysteine (SAH); S-adenosyl homocysteine hydrolase (SAHH); Homocysteine (Hcy); Tetrahydrofolate (THF); 5-methyltetrahydrofolate (5-MTHF); Methionine synthase (MTR); Betaine homocysteine S-methyltransferase (BHMT); Cystathionine β-synthase (CBS); L-cystathionine (L-CYS); Cystathionine γ-lyase (CTH); Glutamate-cysteine ligase (GCL); Glutathione synthetase (GSS); Dimethylglycine (DMG); Glutathione (GSH); Glutathione disulfide (GSSG); Glutathione peroxidase (GPX). *\*p* < 0.05; *\*\*p* <0.01; *\*\*\*p* < 0.001; *ns*: not significant; Mann-Whitney test, a, b (left); two-way ANOVA followed by Bonferroni’s correction, b (center, right), e, g, h; Wilcoxon test, c, d; two-tailed paired *t* test, i.

Using a syngeneic mouse model of induced liver metastasis by intrasplenic injection of Diff or Mes PK4A cells, we also showed that Mes cells produced a higher number of metastatic foci with undifferentiated phenotype than Diff cells (**Figure 2B, Supp Figure 2C**). We further examined the oncogenic features of mesenchymal and differentiated PDA cells and observed that the clonogenic ability (**Figure 2C left**), as well as the potential for individual (**Figure 2D left**) and collective (**Figure 2E**) migration were increased in Mes PK4A cells compared to Diff ones, as it did in Pop2 compared to Pop1 KPC cells (**Figure 2C right, D right**). As expected, EMT induction in TGFβ-treated PK4A and Pop1 KPC cells also increased their migration capacity (**Supp Figure 2D**). Moreover, when cultured under ultra-low attachment conditions, Mes PK4A cells and TGF-β-treated PK4A cells formed larger spheroids and organoids compared to Diff PDA or NT PK4A cells (**Figure 2F, Supp Figure 2E**). These results demonstrate that mesenchymal PDA cells have enhanced proliferative, invasive, and migratory abilities together with a more aggressive behavior *in vivo* compared to differentiated cells.

Since Mes PDA cells present high levels of the methionine cycle and transsulfuration enzymes, we examined if this was correlated with a metabolic advantage compared to their differentiated counterparts. We observed that Mes PDA cells and Pop2 KPC cells showed a higher maximal mitochondrial respiration capacity than Diff PK4A and Pop1 KPC cells, respectively (**Figure 2G, Supp Figure 2F**). As the transsulfuration pathway supplies cysteine to the GSH pathway, which is essential for the antioxidant function of tumor cells, we challenged our PDA cell models with menadione to boost ROS levels and evaluate the ability of tumor cells to balance ROS production. Indeed, ROS levels were significantly higher in Diff PK4A cells, as well as in Pop1 KPC cells, than in the Mes PK4A and Pop2 KPC cells, specifically at high menadione concentration (**Figure 2H, Supp Figure 2G**), indicating that PDA cells with a strong cEMT phenotype have a stronger antioxidant potential and can efficiently counterbalance the high generation of ROS. Next, we broadly investigated the abundance of several metabolites of the methionine cycle and associated pathways in Mes *vs* Diff PK4A cells using targeted metabolomics. We observed that the substrate (SAH) and the product (Hcy) of SAHH were drastically increased in Mes PK4A cells compared to Diff PKA4 cells (**Figure 2I**). Importantly, Mes cells harbored elevated levels of GSH, indicated by the higher GSH/GSSG ratio, and low levels of products of cysteine oxidation (taurine and sulfate) compared to Diff cells, suggesting that the cysteine pool is mainly used for GSH production in Mes cells.

### Targeting SAHH in mesenchymal PDA tumor cells alters their proliferative and antioxidant functions

Given the upregulation of key enzymes of the methionine cycle, transsulfuration, and GSH pathways, and the high levels of Hcy and GSH pools in Mes cells, we hypothesized that the activation of these pathways was required to support Mes cell functions. Therefore, we next examined the role of SAHH, that connects the methionine cycle with the transsulfuration pathway, on the mesenchymal behavior of PDA cells. First, we used DZA (3-Deazaadenosine), a specific chemical inhibitor of SAHH and observed that the IC_50_ of DZA in cell viability was significantly lower for Mes PDA cells than for the Diff ones (99.8 µM vs 241.8 µM) (**Figure 3A**), suggesting that Mes cells were more sensitive to SAHH inhibition. Moreover, increasing DZA concentrations over time drastically limited the proliferation of Mes PK4A (**Figure 3B**), and of KIC-LM cells (**Supp Figure 3A**). Importantly, we showed that DZA treatment induced an arrest of Mes PK4A cell proliferation rather than cell death, as the observed accumulation of CFSE was proportional to the increasing concentration of DZA (**Supp Figure 3B**). These findings were clinically validated on PDA cells derived from three distinct patient-derived xenografts (PDX) of PDA and LM. Interestingly, the three PDA- and paired LM-PDX-derived tumor cells were up to 80% enriched in cEMT cells (**Supp Figure 3C**). Consistently, upon DZA treatment, both PDA- and LM-derived-PDX tumor cells showed a decreased proliferation rate with increasing DZA concentration (**Figure 3C, Supp Figure 3D**). As an alternative strategy to study the role of SAHH, we genetically disrupted *Ahcy* (the gene coding for SAHH) using CRISPR-Cas9 gene editing and generated SAHH-deficient Mes PK4A cells (polyclonal (SAHH-def PC) and clonal (SAHH-def#1 clone) populations, as well as SAHH-deficient LM KIC cells (SAHH-def #A and #B clones) (**Supp Figure 3E**). Downregulation of SAHH induced a significant decrease in the proliferation of both primary and LM mesenchymal cells (**Supp Figure 3F, G**). The impact of SAHH inhibition on mesenchymal cell growth was maintained in cells cultured in 3 dimensions (3D) as DZA treatment drastically inhibited the growth of Mes PK4A and Pop2 KPC cells when cultured as spheroids or organoids, and this effect was validated in patient-derived PDA and LM cells (**Figure 3D, Supp Figure 3H**). Finally, SAHH targeting drastically limited the migration of Mes PK4A cells (**Figure 3E**). Thus, blocking the methionine cycle by both chemical and genetic targeting of SAHH induced the loss of aggressiveness features in mesenchymal PDA cells. Similar results were obtained upon chemical inhibition of the transsulfuration enzyme CTH, which lies downstream of SAHH. Indeed, incubation of Diff and Mes PK4A cells with the specific inhibitor DL-Propargylglycine (DLP) did not impact the proliferation of Diff, but it importantly reduced the proliferation of Mes PK4A cells (**Supp Figure 3I**).

**Figure 3.**
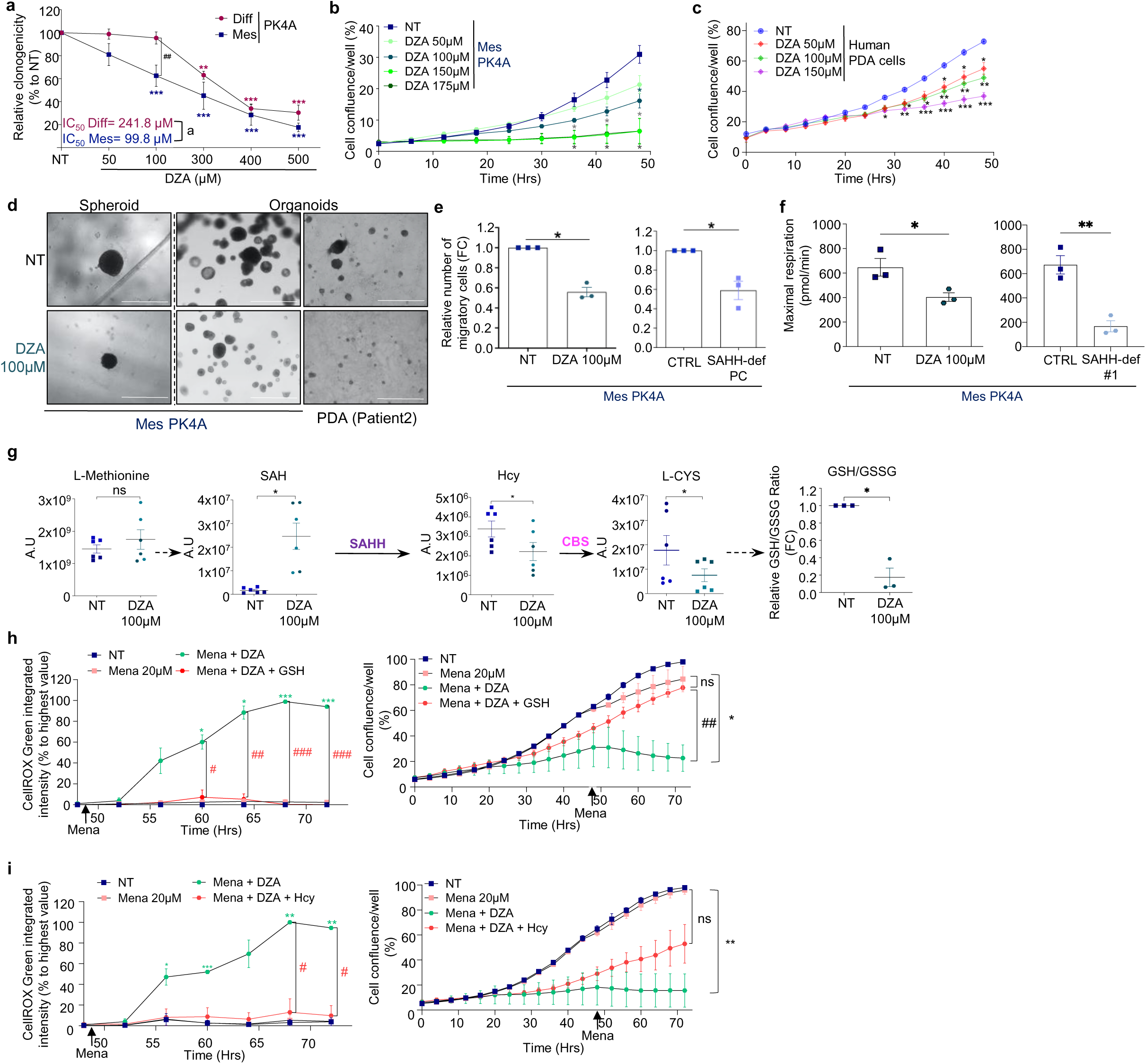
Inhibition of the methionine cycle decreases oncogenic and metabolic abilities of mesenchymal PDA cells. **a,** DZA-induced decrease in the clonogenic capacity of Diff and Mes PK4A throughout 48hrs, expressed by the percent mean±SEM relative to non-treated (NT) control. IC_50_ values indicated for each cell type. * DZA vs respective NT; ^#^ Mes vs Diff for a given DZA concentration. **b,c** Confluence of NT and DZA-treated Mes PK4A cells (**b**) and human PDA-derived primary cells (**c**, n=3 patients) over 50 hrs expressed as mean±SEM. Significant differences illustrated between NT and treated conditions at each time point. **d,** Mes PK4A spheroids (left) and Mes PK4A (center) and human PDA (right) organoids treated with DZA for 7 days (spheroids) or 48hrs (organoids). Representative pictures from 3 independent experiments (spheroids) or 3 technical replicates (organoids). Scale bar, 1 mm. **e,** Migration comparison between NT and DZA-treated Mes PK4A cells (left) or between control (CTRL) and SAHH-deficient Mes PK4A cells (right) over 48h. Values are mean±SEM of the FC relative to NT or CTRL Mes PK4A cells respectively. **f,** Mean±SEM of the maximal respiration capacity of NT or DZA-treated Mes PK4A cells (left) or of CTRL and SAHH-def#1 (clone#1) Mes PK4A cells (right) over 48h. **g,** Levels of intracellular L-Methionine, SAH, Hcy, and L-CYS in NT or DZA-treated Mes PK4A cells over 48hrs. Data expressed in A.U. as mean±SEM from 2 independent experiments with 3 replicates each. The GSH/GSSG ratio measured in NT or DZA-treated Mes PK4A cells is expressed as the mean±SEM of the FC relative to NT. **h and i,** Total ROS production (left) and proliferation (right) in Mes PK4A cells NT or treated with 20 µM menadione (Mena) alone, or combined with 100µM of DZA, or with 100µM of DZA and 1mM of GSH (**h**) or Hcy (**i**) for 72hrs. ROS levels expressed as mean±SEM percentage of the highest value, while confluence is expressed as mean±SEM and significance was defined at the end point. * “Mena 20µM” vs “Mena+DZA”. ^#^ “Mena+DZA” vs “Mena+DZA+GSH/Hcy”. All results are from at least three independent experiments unless specified otherwise. ^a,^ *^, *#*^p < 0.05; *\*\*^, ##^p* <0.01; *\*\*\*^, ###^p* < 0.001; *ns*: not significant. Two-way ANOVA with Dunnett’s correction, a; repeated measures two-way ANOVA with Dunnett’s correction, b, c, h and i (left); unpaired t-test with Welch correction, a (for IC_50_); unpaired t-test, f, h and i (right); paired t-test, g; Wilcoxon test, e, g (GSH/GSSG ratio).

We further examined the metabolic consequences of SAHH inhibition in Mes PK4A cells. Decreased enzymatic activity of SAHH upon DZA treatment or *Ahcy* knock-down (**Supp Figure 3J**) impaired mitochondrial respiration (**Figure 3F**), provoked an accumulation of SAH and decreased the pools of Hcy, L-CYS, and of the GSH/GSSG ratio (**Figure 3G, Supp Figure 3K**). Thus, we further evaluated the influence of low Hcy levels and GSH production upon SAHH inhibition on antioxidant capacity and viability of Mes cells. Chemical SAHH inhibition in mesenchymal cells (*i.e.* Mes PK4A and LM-cells) drastically increased the levels of menadione-induced ROS compared to non-treated or SAHH expressing cells, and this was associated with cell proliferation arrest (**Figure 3H, I, Supp Figure 3L, M**). Importantly, GSH or Hcy addition abrogated menadione-induced ROS production upon SAHH inhibition and partially rescued mesenchymal cell proliferation (**Figure 3H, I, Supp Figure 3L, M**). These results suggest that the methionine cycle, and more specifically SAHH, contributes to the proliferative and antioxidant capacities of mesenchymal cells by sustaining high levels of Hcy and GSH.

### SAHH inhibition in mesenchymal cells disrupts calcium homeostasis and decreases EMT features

To further explore the influence of the acute inhibition of SAHH, we performed RNAseq in non-treated (NT) or DZA-treated Mes PK4A cells cultured for 48h. The analysis showed more than 7,600 differentially expressed genes following DZA treatment (p-value ≤ 0.05, **Supp Table 4**). GSEA enrichment mapping revealed significantly (FDR ≤ 0.01) upregulated nodes (in red) related to mitochondrial and ribosome/RNA metabolism, DNA regulation, as well as downregulated nodes (in blue) including TGFβ signaling and EMT regulation, regulation of calcium (Ca^2+^) homeostasis and transport, endoplasmic reticulum stress (ERS) response, ion homeostasis, lipid and phospholipid metabolism, catabolic and autophagy processes, cell differentiation and cytoskeleton processes, as well as extracellular matrix components and receptors (**Figure 4A, Supp Table 5**). The ER is the primary reservoir of Ca^2+^ and contributes to cytosolic Ca^2+^ homeostasis ^32,33^ and to the ERS response, namely the unfolded-protein response (UPR), that can potentiate the EMT^34^ (**Supp Figure 4A**). Interestingly, many genes involved in Ca^2+^ regulation (*Trpm4, Cacna2d1/1h, Orai3, Atp2c2*), in EMT (*Vimentin, Zeb1, Itgav*) and in ERS response (*Xbp1, Atf4/6, Herpud1, Creb3l1, Ddit3*) were commonly downregulated after DZA treatment, whereas genes involved in Ca^2+^ transport (*Orai2, Ryr2)* and signaling (*Camkk2*) were upregulated (**Figure 4B**). The Ca^2+^ channel ryanodine receptor (RyR) together with the inositol triphosphate receptor (IP_3_R) promote the Ca^2+^ outflux from the ER stores and facilitate Ca^2+^ transfer from the ER to the mitochondria. Hence, the tight regulation of Ca^2+^ homeostasis is performed by proteasome-mediated degradation of RyR and IP_3_R, that is controlled by HERP1 (Homocysteine-inducible ER stress protein, coded by *Herpud*), an ER resident protein (**Supp Figure 4A**). Interestingly, under increased ROS conditions, HERP1 limits the channel-dependent efflux of Ca^2+^ from ER, and protects cells against mitochondrial Ca^2+^ overload and subsequent damages induced by oxidative stress^35^. Upon SAHH inhibition, we observed loss of HERP1, downregulation of the Ca^2+^ regulator TRPM4, and increased levels of CAMKK2, a marker of cytosolic Ca^2+^ overload^36^ (**Figure 4C, Supp Figure 4b**), suggesting that SAHH inhibition disrupts intracellular Ca^2+^ homeostasis by inducing a Ca^2+^ leak in the ER. Consistently, we observed that intracellular Ca^2+^ accumulates over time in DZA-treated Mes PK4A cells (**Figure 4D, Supp Figure 4C**), confirming that under SAHH inactivation, the fine tuning of Ca^2+^ homeostasis was lost. In addition, we observed that disruption of Ca^2+^ homeostasis was associated with downregulation of the ERS pathway. Indeed, despite the increased levels of BIP, an upstream regulator of the UPR pathway, several markers of the ERS were downregulated in SAHH-deficient cells such as the inositol-requiring enzyme 1α (IRE1α), the activating transcription factors 4 and 6 (ATF4 and ATF6), and the C/EBP homologous protein expression (CHOP), an ATF4 target (**Figure 4E right, Supp Figure 4B**), suggesting inefficient ERS response. These results are in line with the transcriptomic data showing the downregulation of the transcription factor X-box binding protein 1 (*Xbp1*, target of IRE1α), and of *Herpud1*, target of XBP1 (**Figure 4B**). We also observed that the loss of Ca^2+^ balance in Mes cells upon SAHH inhibition was accompanied by decreased levels of EMT drivers such as CD51 (ITGAV), CD106, N-CAD, SLUG, and Vimentin (**Figure 4E right, Supp Figure 4B**), as suggested by our RNAseq analysis and by low levels of TRPM4 known to suppress EMT, cell proliferation and invasion^37^.

**Figure 4.**
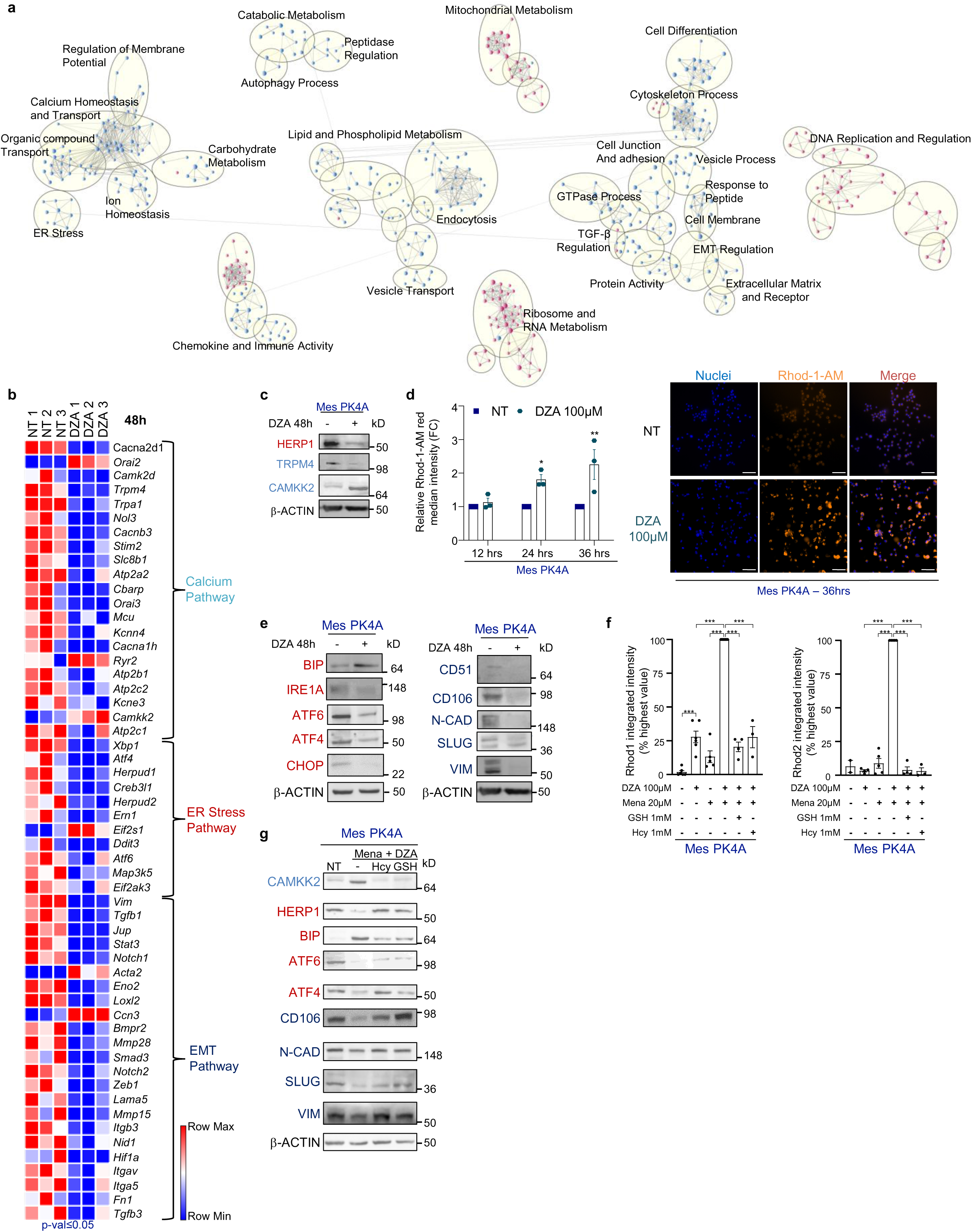
Blocking the methionine cycle induces dysregulation of calcium homeostasis and down-regulation of EMT markers. **a,** Enrichment map representing GSEA of DZA-treated compared to non-treated (NT) Mes PK4A cells over 48hrs. Each node represents a gene set associated with a specific GO (Gene Ontology) term and related nodes cluster into a common biological process (grey circles). Red and blue nodes illustrate respectively up- and down-regulated gene sets in DZA compared to NT. **b,** Heatmap showing the differentially expressed genes in DZA-treated Mes PK4A cells involved in calcium homeostasis, ER stress regulation and EMT pathways. **c,** Representative immunoblot of HERP11, TRPM4 and CAMKK2 in Mes PK4A cells treated or not with 100µM DZA for 48hrs. β-Actin was used as loading control. **d,** Left: X-Rhod1-AM accumulation in NT and DZA-treated Mes PK4A cells for 12, 24, and 36 hrs. DZA-treated cell X-Rhod1-AM intensity is expressed as FC relative to NT for each time point and represented as mean±SEM. *\** NT *vs* DZA-treated cells per time point. Right: Representative images of Rhod-1-AM fluorescence (orange) in NT or DZA-treated Mes PK4A cells for 36hrs. Nuclei were visualized with DAPI (blue). Scale bar, 50µM. **e,** Representative immunoblots of ER stress regulators (left) and EMT markers (right) in Mes PK4A cells treated or not with 100µM DZA for 48 hrs. β-Actin was used as loading control. **f,** X-Rhod1-AM (left) and Rhod2-AM (right) accumulation in Mes PK4A cells control or treated with DZA and Mena alone or combined, and with DZA+Mena combined with GSH or Hcy. Rhod1/2 levels are expressed as mean±SEM percentage of the highest intensity value from 2-11 independent experiments. **g,** Representative immunoblot of CAMKK2, ER stress and EMT markers in NT and Mena+DZA-treated Mes PK4A cells combined or not with GSH or Hcy for 48hrs. β-Actin was used as loading control. All results are from at least three independent experiments unless otherwise specified. \**p* < 0.05; \*\**p* <0.01; \*\*\**p* <0.001. Two-way ANOVA with Dunnett’s correction, d; One-way ANOVA with Tukey’s correction, f.

Finally, to evaluate if the Ca^2+^ imbalance in DZA-treated Mes cells was a consequence of the massive increase in ROS levels, we quantified the Ca^2+^ levels in DZA-treated cells with or without rescue by Hcy or GSH. We observed a strong increase in both cytosolic and mitochondrial Ca^2+^ after treatment with menadione and DZA (**Figure 4F**). This Ca^2+^ overload was associated with increased mitochondrial ROS (**Supp Figure 4D**) in addition to the previously observed high levels of total cell ROS (**Figure 3H, I left panels**). Interestingly, supplementation with GSH or Hcy that impedes ROS production (**Figure 3 H, I, left panels, Supp Figure 4D**) limits Ca^2+^ overflow in both mitochondria and the cytosol (**Figure 4F**), and this rescue was associated with the recovery of levels of CAMKK2, HERP1, ERS and EMT markers close to control levels in mesenchymal cells (**Figure 4G**). On the other hand, the addition of the calcium chelator BAPTA (1,2-bis(o-aminophenoxy) ethane-N,N,N’,N’-tetraacetic acid) was inefficient to blunt the accumulation of Ca^2+^ and the high production of mitochondrial ROS in Mes PK4A cells (**Supp Figure 4E**), suggesting that increased ROS levels in such conditions is the main contributor of the loss of Ca^2+^ homeostasis.

Collectively, these data show that SAHH-deficient mesenchymal PDA cells fail to balance ROS overproduction due to the lack of sufficient levels of GSH and Hcy. This induces intracellular Ca^2+^ overload that diminishes the capacity of these cells to maintain ER homeostasis and the high levels of mesenchymal markers. Thus, mesenchymal cells rely on the activation of the methionine cycle to maintain low ROS levels, Ca^2+^ homeostasis, and their mesenchymal features.

### Inhibition of the methionine cycle suppresses tumor growth and LM formation

Our data show that PDA and LM cells mainly harbor hybrid or cEMT states and that orthotopic or intrasplenic injection of mesenchymal PDA cells leads to aggressive PDA and increased metastatic incidence respectively (**Figure 2**). As mesenchymal PDA cells lose their EMT features upon SAHH inhibition, we examined the effect of pharmacologic or genetic SAHH targeting on mesenchymal PDA growth and LM formation using the two previously described mouse models (**Figure 2**). Both routes of SAHH disruption induced a drastic suppression of mesenchymal PDA growth (**Figure 5A**) in association with a reduced proliferative potential of tumor cells, demonstrated by the weak Ki67 staining in DZA-treated or in SAHH-def tumors, where SAHH inhibition was stable (**Figure 5B, Supp Figure 5A**). Interestingly, the ability of Mes cells to colonize liver lobes and form small to large and undifferentiated metastases was considerably reduced upon DZA treatment or following SAHH knock-down (**Figure 5C, Supp Figure 5B**). Importantly, DZA treatment did not affect mouse body weight (**Supp Figure 5C**). In conclusion, these *in vivo* data establish that inhibition of the methionine cycle, via SAHH targeting, drastically reduces the growth of mesenchymal PDA as well as of metastasis formation.

**Figure 5.**
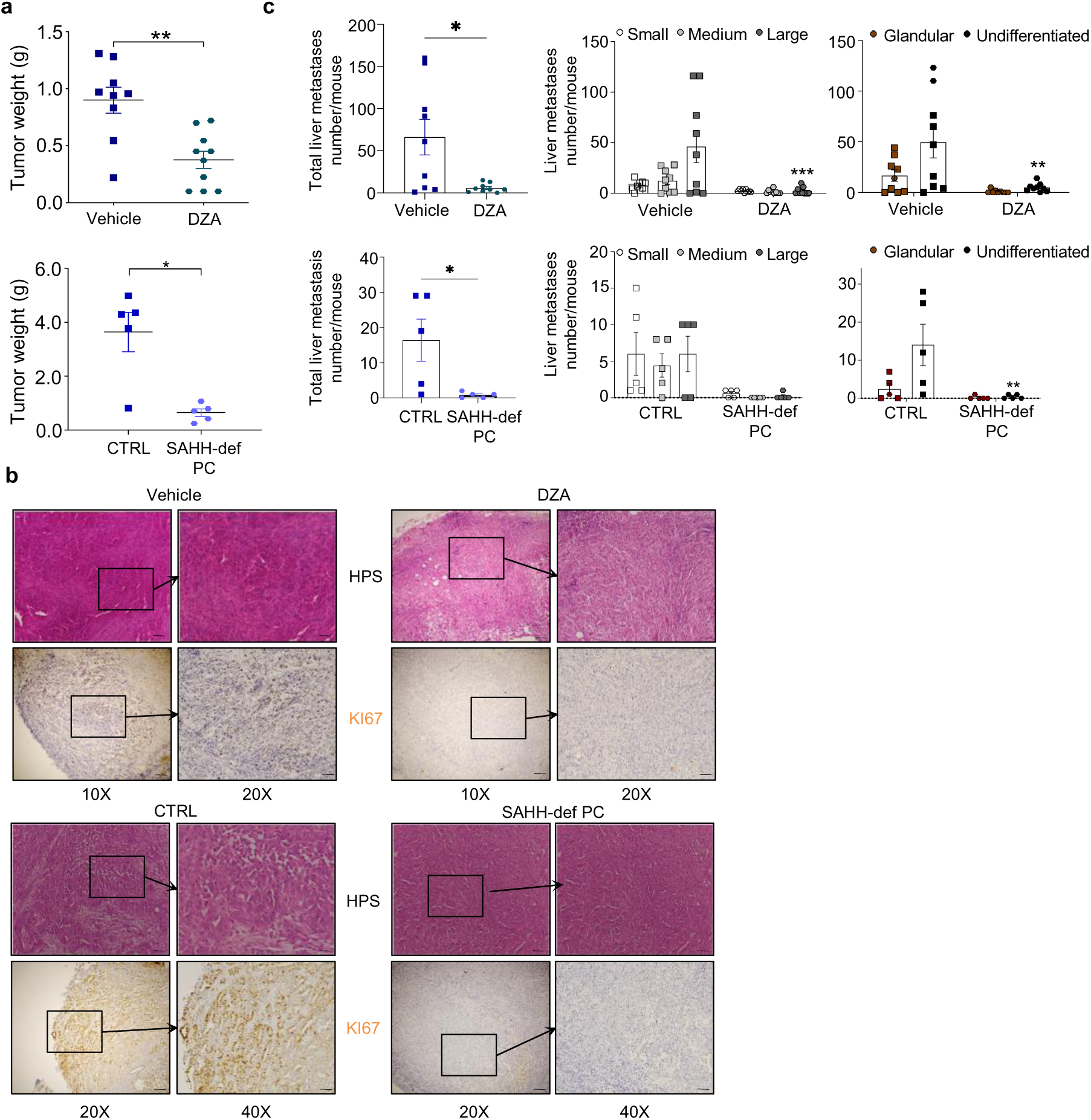
Inhibition of methionine cycle impedes tumor growth and metastasis formation. **a,** Mean±SEM of the weight of the primary tumor of KI mice orthotopically implanted either with Mes PK4A cells (top) and treated with vehicle (n=9 mice) and DZA (20 mg/kg/mice, n=10 mice), or implanted with CTRL and SAHH-def PC Mes PK4A cells (bottom, n=5 mice/experimental group). *\*p* < 0.05; *\*\*p* < 0.01, Mann-Whitney test. **b,** Representative images of HPS staining and KI67 immunostaining of Mes PK4A tumors of KI mice at the end of the treatment (top, n=3 independent experiments) and of CTRL or SAHH-def PC Mes PK4A tumors of KI mice 28 days post-implantation (bottom, n=3 independent experiments). Scale bars: 10X, 100 µM; 20X, 50 μm. **c,** Number of metastases per mouse (1 dot = 1 mouse) counted after implantation of Mes PK4A cells treated with vehicle or with DZA (top, n=9 mice/experimental group), or with CTRL or SAHH-def PC Mes PK4A cells (bottom, n=5 mice each/experimental group) and classified by the total number of metastasis (left), their size (center; small, white dots; medium, grey dots; large, black dots), or their histological status (right; glandular, brown dots; undifferentiated, black dots). Data expressed as mean±SEM. *\*p* < 0.05; *\*\*p* < 0.01; *\*\*\*p* < 0.001. Mann-Whitney test for the total number of metastases, and two-way ANOVA with Bonferroni’s correction for size and pathological status.

### High expression of enzymes of the methionine cycle in human mesenchymal PDA is associated with poor prognosis

To study the clinical relevance of the activation of the methionine cycle in human clinical samples of PDA and LM, we analyzed gene expression data from PDA primary tumors and LM collected from 17 publicly available whole-genome mRNA expression data sets (**Supp Table 6**). The clinicopathological characteristics of the PDA are summarized in **Supp Table 7**. Interestingly, the expression of an EMT expression signature based on *CDH2* and *ITGAV* was higher in LM samples than in PDA samples (p=8.21E-06, Fisher’s exact test; **Figure 6A left**). EMT enrichment in LM was confirmed by analysis of Collisson’s molecular subtypes: 55% of LM *versus* 25% of PDA samples were of “quasimesenchymal” subtype (p=1.16E-05, Fisher’s exact test; **Figure 6A right**). We next performed metagene analysis on the expression of 21 genes involved in the methionine cycle and 24 genes of the transsulfuration pathway (**Supp Table 8**). We observed increased metagene scores for both pathways (**Figure 6B)**, and higher *AHCY* expression (**Figure 6C**) in LM compared to PDA. Importantly, the expression of the mesenchymal marker CD51 and of SAHH was confirmed in tumor cells in clinical PDA and LM samples (**Figure 6D**). Therefore, we assessed the potential prognostic value of *AHCY* and mesenchymal gene expression in the 755 patients informative for overall survival (OS). With a median follow-up of 17 months (range, 1 to 156), the 2-year OS was 38% (95%CI 35-42). We found that the EMT signature correlated with OS (p=1.08E-02, Wald test) as the 2-year OS was 34% in the EMT-high group *versus* 39% in the EMT-low group (p=1.08E-02, log-rank test; **Figure 6E**). Furthermore, *AHCY* expression did not correlate with OS (p=0.85, Wald test, **Supp Table 9**). Interestingly, a prognostic interaction existed between EMT signature and *AHCY* expression. Indeed, in the EMT-high group (CDH2-high and ITGAV-high), high *AHCY* expression was significantly associated with shorter OS (p=2.13E-02, log-rank test, **Figure 6F, left**), whereas in the EMT-low group (CDH2-low and/or ITGAV-low), *AHCY* expression was not associated with survival (p=0.50, log-rank test; **Figure 6F, right**). Collectively, these clinical data show that high levels of *AHCY* expression in mesenchymal PDA are associated with poor prognosis.

**Figure 6.**
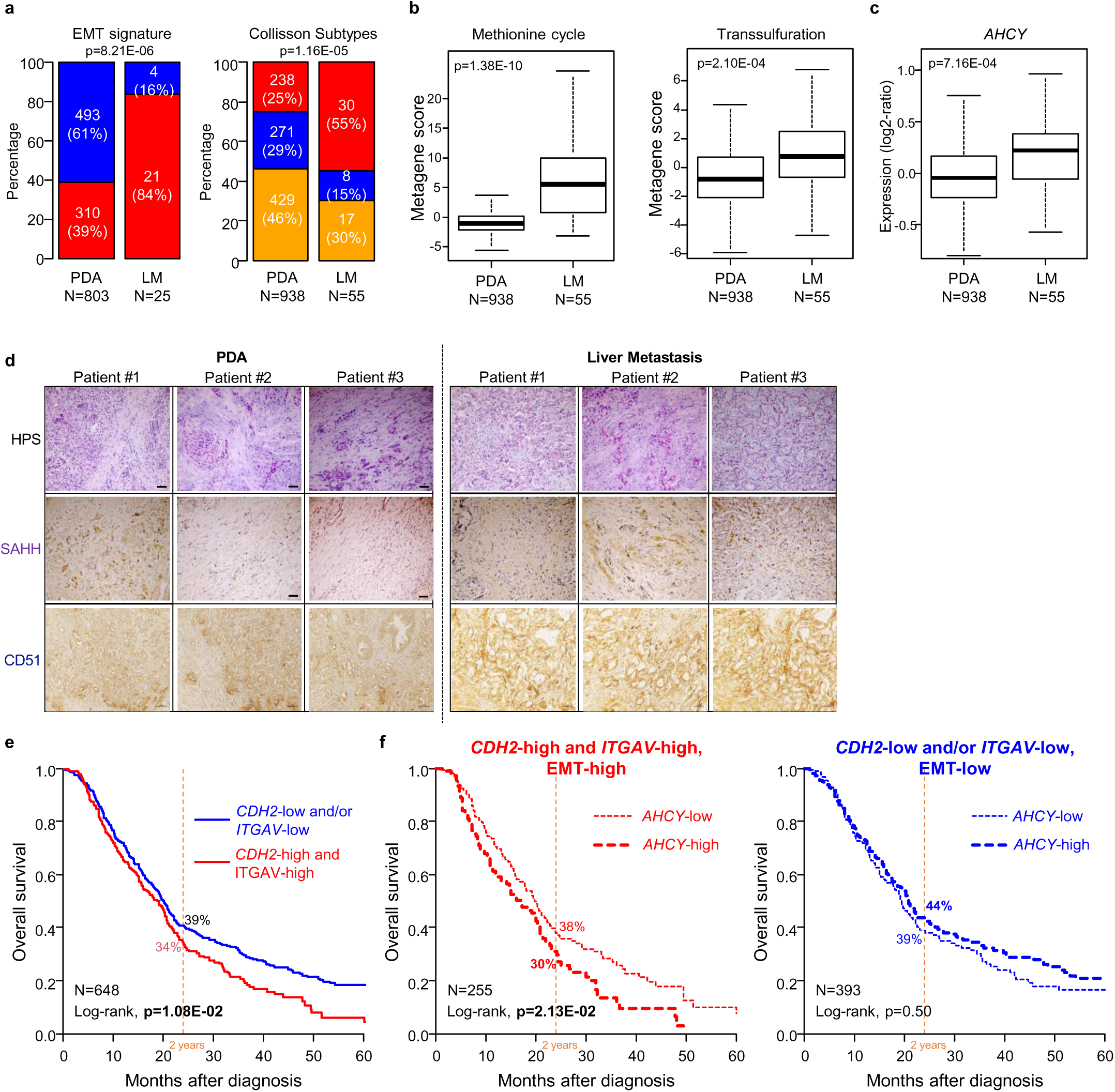
Upregulation of metabolic genes of the methionine cycle in human mesenchymal PDA is associated with poor overall survival. **a**, Bar plot showing the proportion of tissue types categorized by EMT classes (CDH2-low and/or ITGAV-low in blue; CDH2-high and ITGAV-high in red, left) and by Collisson’s molecular subtypes (quasi-mesenchymal, red; exocrine-like, blue; classical, orange; right). **b and c,** Box-and-whisker plots of the expression scores of the methionine cycle and transsulfuration pathway metagenes defined as first component of PCA of genes (b), and of *AHCY* expression (log2-ratio) in PDA (PDA, n=938) and liver metastases (LM, n=55) (c). Student’s t test. **d**, Representative images of HPS staining and of SAHH and CD51 immunostaining in PDA and LM from 3 patients (n=3 independent experiments). Scale bar, 50 µM. **e**, Kaplan-Meier overall survival (OS) curve in the whole population stratified by EMT groups (EMT-low: CDH2-low and/or ITGAV-low in blue; EMT-high: CDH2-high and ITGAV-high in red). **f**, Kaplan-Meier OS curve based on *AHCY* expression (*AHCY*-low in dotted line and AHCY-high in bold, dotted line) within the EMT-high PDA group (red, left) and within the EMT-low PDA group (blue, right).

## DISCUSSION

In this study we explored the activated metabolic pathways that support the mesenchymal phenotype of PDA cells and their contribution to the formation of metastasis. PDA cells subjected to EMT evolve through a continuum of intermediate states that are tightly regulated by EMT-specific transcription factors (TFs). However, pharmacologic targeting of key EMT TFs remains challenging. Here, our data provide insight into the identification and targeting of metabolic regulators of EMT in mesenchymal PDA and in LM that may be used as a promising therapeutic approach to revert or reduce EMT-controlled PDA progression and metastasis. Using *in vitro* and *in vivo* models of metastatic PDA, we show that PDA cells display a plastic hEMT state that can evolve either toward a cEMT or, to a lesser extent, to an epithelial state once tumor cells are established in the metastatic niche. Importantly, we also demonstrate that PDA cells at epithelial and cEMT states can shift toward an hEMT state after cell passaging *in vitro*. This high degree of EMT plasticity confers varying proliferative, invasive, and migratory capacities to PDA tumor cells that depend on the combination of expression of several EMT markers. The presence of multiple effectors that control EMT complicates its targeting or the inhibition of EMT-associated functions. Simeonov et al., showed a continuum in the expression of epithelial and mesenchymal markers in PDA tumor cells in hEMT state, giving rise to a sub-cluster of four distinct hEMT states, from early to late hEMT. The latter state was shown to be the most aggressive and displayed specific proliferative and metabolic capacities^20^. Our data show that PDA cells at low passages co-express epithelial and mesenchymal markers such as E-CAD, CK19 and N-CAD at various degrees. Interestingly, while these cells progress in a cEMT phenotype, N-CAD, commonly assigned to cEMT, shows a slight increase, compared to the drastically increased expression of SNAIL and/or SLUG. This demonstrates that N-CAD is not sufficient to define EMT states even at their extremes.

Interestingly, the same study by Simeonov et al. showed that transition of cells from early to late hEMT state is accompanied by a metabolic change from an OXPHOS to a glycolytic profile, based on the expression of specific metabolic gene sets. This highlights that the metabolic profile of tumor cells is linked to the degree of EMT, with specific metabolic requirements associated with each EMT state^20^. Our work highlights that PDA cells that acquire a cEMT profile over a hEMT or an epithelial state (PDA vs LM, pop1 vs pop2 cells, Diff vs Mes cells, and NT vs TGFβ–treated cells) have increased levels of enzymes involved in the methionine cycle, the transsulfuration and glutathione pathways. Methionine is an essential amino acid that can be regenerated via the re-methylation of Hcy either by MTR using 5-methyl-THF, or by BHMT using betaine as methyl group donor. Recycling of Hcy to provide methionine pool is minimal in PDA cells in which Hcy contributes only up to 10% of methionine production^38,39^.

Consequently, more Hcy is available to produce cysteine to fuel the GSH pathway. Indeed, activated methionine cycle has been previously described in cancer cell lines to provide Hcy that in turn enters the transsulfuration pathway, supporting the notion that Hcy derived from the methionine cycle can contribute to the GSH pool^40^. In our study, the increased levels of SAHH, CTH and GCLC in Mes cells compared to Diff cells, together with high levels of SAH, Hcy and low levels of taurine produced through cysteine oxidation, suggest that cEMT PDA cells rewire Hcy to preferentially fuel the transsulfuration and GSH pathways. These events ultimately contribute to increased levels of GSH. Interestingly, in colorectal cancer (CRC) cells from an APC-deficient, KRAS-mutated mouse model, APC is sufficient to drive expression of SAHH and to increase levels of SAM and SAH. The targeting of SAHH both in organoids and *in vivo* reduces the growth of CRC cells^41^. Similar to CRC cells, the increased sensitivity of Mes PDA cells to the pharmacological inhibition of SAHH compared to Diff cells highlights their dependence on SAHH. Interestingly, decreased activity of SAHH and the resulting low levels of Hcy and GSH in Mes cells are not associated with cell death, but with their ability to proliferate, migrate, and expand in 3D culture. In addition, SAHH targeting in Mes cells increases the levels of menadione-induced ROS beyond the tolerance threshold of the cells, resulting in cell proliferation arrest. This is due to the lack of sufficient Hcy levels to fuel the transsulfuration pathway, resulting in low levels of the antioxidant GSH pools. Indeed, addition of exogenous Hcy and GSH restores the protection against ROS and rescues Mes cell proliferation.

We also found that the blockade of SAHH strongly affects Ca^2+^ homeostasis and the UPR in mesenchymal PDA cells. Mitochondria-associated ER membranes (MAM) are subcellular structures, composed of redox-sensitive calcium handling proteins, that connect ER and mitochondria and allow Ca^2+^ flux between both organelles. In conditions of unmitigated high ROS levels, the tightly controlled Ca^2+^ transfer from ER to mitochondria through MAM is lost, leading to mitochondrial Ca^2+^ escape and release of cytosolic Ca^2+^. Upon DZA we observed low levels of IRE1α, an ER stress inducer that localizes at MAM sites where it interacts with IP_3_R^42^ to control Ca^2+^ flux from ER to mitochondria. Decreased levels of IRE1α together with low levels of HERP1 may contribute to the defective control of intracellular Ca^2+^ transport. Upon DZA, we also observed high levels of BIP, which has a dual role as gate keeper of ER homeostasis upstream of the UPR pathway and as co-chaperone to regulate Ca^2+^ homeostasis. The inability of DZA-treated cells to efficiently induce an ERS response suggests that BIP induction is an attempt to compensate for the Ca^2+^ release into cytosol, rather than to induce an efficient ERS response. Finally, we show that the massive ROS production together with uncontrolled Ca^2+^ cytosolic flux following SAHH inhibition induces a drastic reduction of EMT markers in Mes cells. In these conditions of high oxidative stress, the addition of GSH and Hcy to SAHH-deficient Mes PK4A cells restores Ca^2+^ to basal levels by bringing down ROS, suggesting that the high production of ROS alters Ca^2+^ homeostasis. Rescuing ROS balance and Ca^2+^ homeostasis by addition of Hcy or GSH restores the levels of CAMKK2, HERP, ERS markers and importantly of EMT markers to basal levels observed in Mes PK4A cells. The Ca^2+^ signaling contribution to EMT in cancer cells^43^ was previously illustrated by the TRPM4 channel which regulates intracellular Ca^2+^ flux and promotes β-catenin signaling, ultimately activating EMT and cancer cell migration.^37^ Collectively, our data highlight that the methionine cycle regulates the ROS-Ca^2+^-EMT axis in mesenchymal PDA cells, thereby showing that a metabolic targeting strategy, by inactivation of SAHH, inhibits the EMT features and aggressiveness of PDA cells.

Importantly, genetic or pharmacologic targeting of SAHH *in vivo* was well tolerated in PDA- and LM-bearing mice, and the resulting inhibition of the methionine cycle drastically reduced tumor growth and metastatic burden. Therefore, from a clinical point of view, the use of SAHH as a target for the blocking of the methionine cycle to treat metastatic PDA patients may be an attractive strategy to limit the use of methionine-depleted diets. Although such diets contribute to chemo- and radiotherapy sensitivity^44^, they may affect the already compromised general state of patients with metastatic cancer. Interestingly, patients with a predominantly mesenchymal PDA and high expression of *SAHH* show a low overall survival. This observation suggests that the inhibition of *SAHH*, similar to the abrogation of mesenchymal features of tumor cells, could directly correlate with the improvement of survival of patients with metastatic PDA.

## Methods

### Spontaneous PDA mouse models

Male and female PDA-bearing mice Pdx1-Cre; LSL-Kras^G12D/+^; Ink4a/Arf^fl/fl^ (KIC) and control LSL-Kras^G12D^; Ink4a/Arf^fl/fl^ (KI) littermates were obtained as previously described^45^. Male and female PDA-bearing mice Pdx-1-Cre; LSL-Kras^G12D/+^; LSL-Trp53^R172H/+^ (KPC) were obtained by breeding Pdx-1-Cre; with LSL-Kras^G12D/+^;LSL-Trp53^R172H/+^ mice^46^. All animal care and experimental procedures were performed in agreement with the Animal Ethics Committee of Marseille under reference (APAFIS#A1301303-2019020813566398). After sacrifice at 9 weeks (KIC mice) or between 6 to 8 months (KPC mice), pieces of primary tumor or liver metastasis were either fixed in 4% (wt/vol) formaldehyde, dissociated for tumor cells isolation, or snap frozen in cold isopentane for further analysis.

### Laser Capture microdissection of primary tumor and liver metastasis of KIC mice

Laser capture microdissection (LCM) was performed at the microdissection laboratory of the PRIMACEN platform, University of Rouen (France) as previously described^47^. Livers and PDA (n=4 KIC primary tumor and matched liver metastasis) were fixed in dry-ice–cooled isopentane, then sectioned (12–14 μm sections) on a cryostat (Leica CM1950; Leica, Rueil-Malmaison, France), and transferred to PEN-coated (poly-ethylene-naphthalene; Membrane Slide 1.0 PEN; Carl Zeiss, Le Pecq, France) glass slides, previously treated with RNAse Zap (Sigma-Aldrich, St Louis, MO, USA) and UV-activated. The sections were fixed in 70% ethanol for 10 s and stained with haematoxylin and eosin (H&E). Briefly, the sections were immersed in modified Harris haematoxylin solution (Sigma-Aldrich) for 1 min, and rinsed in 100% ethanol before staining in Eosin Y (Sigma-Aldrich) for 30 s. The samples were dehydrated for a few seconds in increasing ethanol concentrations (70-100%). LCM of PDA and LM tumor cells was performed with a Palm Robot MicroBeam apparatus (Carl Zeiss), by applying a pulsed UVA laser beam (focal spot <1 μm) through an inverted microscope with a 20X objective. Dissection of metastatic tumor cells at the edge of the contact with the liver parenchyma was avoided. PDA and LM areas were collected by catapulting into the cap of a tube containing 34 μl of RLT lysis buffer (Qiagen, Courtaboeuf, France) with 1% β-mercaptoethanol (Sigma-Aldrich).

### RNA extraction and purification

Total RNA was extracted using RNeasy Micro kit (Qiagen) in accordance with the manufacturer’s instructions. The RNA was eluted with 10 μl of RNase-free water. DNase I treatment (RNase-Free DNase Set; Qiagen) was performed to remove contaminating genomic DNA. The quality of the RNA was assessed, and the quantity was estimated by microfluidic capillary electrophoresis (Agilent 2100 Bioanalyzer; Agilent Technologies, Massy, France).

### Tumor cell isolation and cell culture

KIC or KPC derived PDA and LM were cut in small pieces, submitted to mechanic dissociation and digestion using the gentleMACS Octo Dissociator (Miltenyi) according to Miltenyi protocol, GentleMACS C Tube, and the tumor dissociation kit (Miltenyi) according to supplier’s protocol. After dissociation, KIC-or KPC-derived PDA and LM cell pellets were filtered in 100 µM cell strainer. Human PDA (PT) and liver metastasis (LM)-derived primary cell cultures were obtained as previously described^48,49^. PK4A1 and 2 cell lines were isolated from PDA of KIC 1 and 2 mouse, respectively, as previously described^45^, and have been used for each experiment to limit tumor batch effect. For cell passaging, PK4A1 and PK4A2 cells were detached from plates with diluted accutase (50% PBS, 5mM EDTA) and seeded at early passages (passages 12-19, Diff), intermediate passages (passages 20-25) until late passages (passages 30-40, Mes). EpCam^+^ subset-derived Pop1 and cEMT subset-derived Pop 2 of KIC1, KIC2, KPC1 and KPC2 were characterized as described in « Flow cytometry analysis for epithelial and mesenchymal cell phenotyping” section. All cells were maintained in complete medium: 25mM glucose, glutamine free Dulbecco’s modified Eagle’s medium (DMEM) without pyruvate, supplemented with 2mM of L-glutamine, 10% (vol/vol) FCS. Cells were cultured with 1% antibiotic/antimycotic solution at 37°C, 21% oxygen in a 5% CO_2_ incubator and were mycoplasma free. For TGF-β stimulation, PK4A at early passages 14-19 or KPC Pop1 cells were plated and next day were treated with 5 ng/mL of TGF-β for indicated time. For DZA (Sigma Aldrich, #D8296) stimulation, PK4A cells were plated in multi-well plates. Twenty-four hours after plating, cells were treated with indicated concentrations or 100µM of DZA for indicated time. The IC_50_ was calculated using a non-linear regression with GraphPad software. For GSH stimulation, PK4A cells were plated in multi-well plates. Twenty-four hours after plating, cells were treated with 1mM of GSH for indicated time in a new media culture. For L-Homocysteine stimulation, PK4A cells were plated in multi-well plates. Twenty-four hours after plating, cells were treated with 1mM of L-Homocysteine for indicated time in a new media culture.

### Fluorescence-activated cell sorting (FACS) and flow cytometry

After 2 to 3 passages following their isolation, PDA and LM tumor cells were detached from plates with diluted accutase, rinsed once with PBS and either examined (see section “Flow cytometry analysis for epithelial and mesenchymal cell phenotyping” below) or cell sorted based on their surface levels of epithelial and mesenchymal markers. For fluorescence-activated cell sorting (FACS), PDA cells were incubated 15-20 min with indicated labelled antibodies at 4°C in the dark. Cells were rinsed with PBS and 500,000 cells/500 µL buffer (PBS, BSA 5%, EDTA 5 mM) were sorted and analyzed on a cell sorter (FACSAria II cell sorter or FACSMelody cell sorter, BD), and further analyzed with FlowJo, to select Epcam^+^ cells and cEMT cells subsets. To select those 2 subsets, Sytox Green (2 drops/tube) (ThermoFisher Scientific) was used to exclude dead cells and select only Sytox green negative alive cells. Then, tumor cells with positive double staining (CD24^+^/CD44^+^ cancer cell markers^6^) and negative staining for CD90 (*i.e.* fibroblast marker) were selected (CD90^-^/CD24^+/^CD44^+^). Tumor cells were then gated first with EpCam marker, and CD90^-^/CD24^+^/CD44^+^/EpCam^+^ cells were sorted as Epcam^+^ cells. EpCam^-^cells were then gated according to CD51 marker, independently of CD106 expression (CD106^+/-^). CD90^-^/CD24^+^/CD44^+^/EpCam^-^/ CD106^+/-^/CD51^+^ cells were sorted as cEMT cells (Supplementary Fig. 1D, F). After 2 to 3 passages EpCam^+^ and cEMT cells were further characterized for epithelial and mesenchymal marker levels at their cell surface, as described below.

### Flow cytometry analysis for epithelial and mesenchymal cell phenotyping

To examine the epithelial and mesenchymal phenotype of PDA and LM of KIC1, 2, 3, and KPC1 (Figure 1F), and of EpCam^+^ subset-derived Pop1 and cEMT subset-derived Pop 2 of KIC1, KIC2, KPC1 and KPC2 (Supp Figure 1G), as well as of PK4A1 and PK4A2 Diff and Mes cells (Supp Figure 1I), cells were plated at 150 000 cells in 12 wells plate. After 48hrs, cells were detached from plates with diluted accutase, rinsed once with PBS and incubated 15-20 min with indicated labelled antibodies at 4°C in the dark. Cells were rinsed with PBS and resuspended in (PBS, BSA 5%, EDTA 5 mM) buffer and analyzed by flow cytometry (LSR Fortessa cell analyzer, BD) followed by Flow-Jo analysis. To determine the percentage of epithelial, hEMT and cEMT cells in each population, alive cancer cells were selected by negative staining of Sytox green (ThermoFisher Scientific) and then gated according to their expression of EpCam marker. EpCam^+^ cells were quantified as EpCam^+^ subset. EpCam^-^population was gated according to its expression of CD51 marker. EpCam^-^/CD51^+^ cells, either CD106 positive or negative (CD106^+/-^), were identified as cEMT cells. Finally, EpCam^-^/CD51^-^cells were quantified as hEMT cells (Supp Figure 6).

### Murine PDA amino acid metabolism transcriptome analysis

All expression data come from 4 pairs of primary tumors (PDA) and matched liver metastasis removed from 9-week-old KIC mice. Gene expression was analyzed by Mouse Gene 2.0 ST Array (Affymetrix). Bio-informatic analysis was performed in R 3.5.1 environment. We focused the differential expression analysis on the 409 genes associated to the amino acid metabolism that we identified based on KEGG pathways. From these 409 genes, Significance Analysis of Microarray (SAM) was performed on the 162 most variable genes between primary tumors and matched liver metastasis pairs (Standard deviation ≥ 0.5), using the function *sam* with default settings of the R package *siggenes* available on Bioconductor. SAM identified 81 significant Differentially Expressed Genes (DEGs) (FDR = 0.16), all upregulated in metastases. Using Ingenuity Pathway Analysis (IPA), enrichment was performed on these 81 amino acid genes and 11 metabolic pathways were found significantly upregulated (p_value_ ≤ 0.05).

### Protein extracts and western blots

PDA and LM tumor cells from KIC1 (Figure 1D, 1G) and KPC1 mice (Figure 1G) and PK4A1 and PK4A2 cells (Figure 1H, 4C, F, G) were lysed in 50 mM Tris pH 8, 150 mM NaCl, 1% NP-40, 0,5% proteases inhibitors cocktail, prior to centrifugation (14,000 rpm, 10 min, 4°C). Supernatants were collected, and protein concentration was evaluated using Bradford protocol. Whole cell protein extracts (20-50 µg/lane) were resolved by SDS/PAGE using a 10% (vol/vol) acrylamide gel and transferred onto 0.2 µm nitrocellulose membranes. After a blocking step (BSA Buffer), primary antibody was incubated O/N at 4°C (in BSA Buffer). The following day, the membranes were rinsed 3 times with TBST (Tris Buffered Saline with Tween20) and then incubated 1 hr at room temperature with secondary antibody (in BSA buffer). ECL protein detection was performed with a Fusion Fx7 chemiluminescent imager and the quantification of the amounts of proteins of interest was determined by densitometry using Image J software (NIH) and normalized to β-actin or β-tubulin or to total loaded protein stained with Ponceau Red.

### Immuno-histochemistry

Formalin-fixed paraffin-embedded, 5 µm-thick human or mouse organs sections were deparaffinized in xylene and rehydrated through a series of graded ethanol concentrations. Antigen retrieval was performed in citrate buffer pH6/Tween 0.05% or target retrieval solution at pH 6 or 9, before the quenching of endogenous peroxidase activity (3% (vol/vol) H_2_O_2_). Tissue sections were then incubated with indicated primary antibody. Immunoreactivity was visualized using the Vectastain ABC kit or with biotinylated goat anti-rabbit, goat anti-mouse, goat anti-rat antibodies and peroxidase-conjugated streptavidin from Dako. Peroxydase activity was revealed using liquid DAB+ substrate chromogen system. Counter-staining in Mayer’s Haematoxilin was followed by a bluing step in sodium bicarbonate buffer (0.1% in water) before final dehydration and mounting of the sections.

### Murine models of induced PDA and liver metastasis *in vivo*

Under isoflurane anesthesia (induction: 4% (vol/vol) and maintenance: 2% (vol/vol)), mice were injected subcutaneously with 0.2 mg/kg buprenorphine (Vetergesic; Sogeval) and lidocaine (Xylovet; Ceva) was administered at 3.5 mg/kg by infiltration in the abdominal cavity.

#### Induced PDA in syngeneic mice

9 to 12 weeks-old male and female KI mice were orthotopically grafted with 1.10^6^ of either Diff or Mes PK4A cells (n=6 mice each), or of CTRL Mes PK4A or SAHH-def PC Mes cells (n= 5 mice each). At the time of sacrifice (day 28), tumors were removed, weighted and fixed in 4% (wt/vol) formaldehyde. For DZA-treated mice, 9 to 12 weeks-old male and female KI mice were orthotopically implanted with 1.10^6^ of Mes PK4A cells and treated at day 7 post implantation with vehicle (n=9 mice) or 20 mg/kg/mice of DZA (n=10 mice) twice a week for 3 weeks. At the time of sacrifice (day 28), tumors were removed, weighted and fixed in 4% (wt/vol) formaldehyde.

#### Induced liver metastasis in syngeneic mice

50 000 of Diff or Mes PK4A cells (n=5 mice each), and of CTRL Mes PK4A or SAHH-def PC Mes cells (n=5 mice each) cells were injected in the spleen of 9-to-12-week-old syngeneic KI mice. At the time of sacrifice (day 21), livers were removed, weighted and fixed in 4% (wt/vol) formaldehyde. For DZA treatment, 9 to 12 weeks-old male and female KI mice were injected in the spleen with 50 000 of Mes PK4A cells and treated at day 7 post implantation with vehicle or 20 mg/kg/mice of DZA (n=9 mice each), twice a week for 2 weeks. At the time of sacrifice (day 21), livers were removed, weighted, fixed in 4% (wt/vol) formaldehyde. Same region and number of liver lobes were embedded in paraffin in a same cassette. 5μm tissue sections were stained by H&E. H&E-stained slides were scanned at 10X and 20X magnification. Liver metastases characterization was performed by counting the number of metastases per slide (at least n=2 slide per mouse, with same number of lobes for all the samples on one slide per mouse) and then classified into 3 different subtypes: small-, medium- and large-size metastases. In addition, anatomopathological status (undifferentiated and glandular) of metastases was determined for each liver lobe with metastases.

### Clonogenic assays

For clonogenic assays, 4 000 cells were seeded in a 12-well plate and cultured in complete medium. For DZA treatment, 24hrs after seeding, indicated concentrations of DZA was added in each well. Three or two (for DZA treatment experiments) days later, colonies were fixed in 4% (wt/vol) formaldehyde and stained with 0.5% Crystal violet. Crystal violet was discolored with 10% Acetic acid and optic density was measured at 600 nm.

### Migration assays

For migratory assays, 60 000 cells were seeded in a 24-well insert in complete medium with 1% FCS and placed in a 24-well plate containing complete media with 10% FCS. 12hrs after seeding, inserts were fixed in 4% (wt/vol) formaldehyde and stained with 0.5% Crystal violet. Crystal violet was discolored with 10% Acetic acid and optic density was measured at 600 nm.

For DZA treatments, cells were plated at 100 000 cells in 6-well plate in complete medium before treatment. The next day cells were treated with indicated concentrations of DZA. After 48hrs, cells were removed with diluted accutase, counted with blue trypan and 60 000 cells were seeded in a 24-well insert in complete medium with 1% FCS with or without DZA, and placed in a 24-well plate containing complete media with 10% FCS. 12hrs after seeding, inserts were fixed in 4% (wt/vol) formaldehyde and stained with 0.5% Crystal violet. Crystal violet was discolored with 10% Acetic acid and optic density was measured at 600 nm.

For TGF-β treatments, cells were plated at 100 000 cells in 6-well plate in complete medium before treatment. The next day cells were treated with 5 ng/µL of TGF-β. After 48hrs, cells were removed with diluted accutase, counted with blue trypan and 60 000 cells were seeded in a 24-well insert in complete medium with 1% FCS with or without TGF-β, and placed in a 24-well plate containing complete media with 10% FCS. 12hrs after seeding, inserts were fixed in 4% (wt/vol) formaldehyde and stained with 0.5% Crystal violet. Crystal violet was discolored with 10% Acetic acid and optic density was measured at 600 nm.

### Wound healing assays

One thousand cells were seeded in 12-well plates. To inhibit proliferation, cells were rendered quiescent using mitomycin C (0.5µg/mL) 2hrs before scratch. After the scratch, wells were washed with PBS 1X, and fresh medium containing mitomycin C (0.5µg/mL) was added. Images were captured at 0hr and 36hrs after scratch, at 2X magnification. Scratched area recovered by cells was measured using ImageJ software and expressed as percentage of measured initial scratched area at day 0.

### Spheroid assays

One thousand cells were seeded in a 96-well ultra-low attachment plate (Corning) in 200µl of complete medium and cultured in 3 dimensions for 15 days. Medium was replenished twice a week with or without DZA or TGF-β. Pictures of spheroids were captured using Evos microscope at 2, 4, or 10X magnification.

### Organoid assays

Ten thousand cells were seeded in a 48-well plate pre-coated with Matrigel as previously described^50^, in 500µl of organoid medium, and cultured in 3 dimensions for 72hrs. After 24hrs, in indicated experimentations, DZA was added for 48hrs. Pictures of organoids were captured using Evos microscope at 2, 4, or 10X magnification.

### Total ROS quantification

For Rev. Figure 3H, I and rev. Supp 3 K, L fifty thousand cells were seeded in 12-well plate in 1 mL of complete medium. 3 days later, Menadione was added for 1hr and then 2.5 µM of green CellROX probe was added for 30 min at 37°C. All the cells were removed with diluted accutase and were rinsed with PBS and centrifuged (2000 rpm, 5min). Cells pellets were resuspended in FACS buffer (PBS, 5% BSA, 5 mM EDTA) and analyzed by flow cytometry with corresponding lasers (MACSQuant VYB flow cytometer, Miltenyi).

### Oxygen consumption rate (OCR)

Twelve thousand cells were seeded into Seahorse 24-well plate in 1 mL of complete medium. For non-treated cells, 72hrs after seeding, medium was changed with DMEM MitoStress medium supplemented with 10 mM of glucose, 1 mM of Pyruvate and 2 mM of L-Glutamine. For DZA-treated cells, mesenchymal cells were treated for 48hrs with DZA and then medium was changed with DMEM MitoStress medium supplemented with 10 mM of glucose, 1 mM of Pyruvate and 2 mM of L-Glutamine. Cells were analyzed by Mitostress test program from Seahorse software and stimulated with 1µM of Oligomycine, 0,5µM of FCCP and 0,5µM of rotenone/Antimycine. OCR was measured using the Seahorse XF24 Extracellular Flux Analyzer (Seahorse Bioscience, North Billerica, MA, USA).

### Metabolomics

For metabolomic analysis 1.10^6^ cells were seeded into 100 mm dishes in 10 mL of complete medium. For experimentation with DZA, cells were treated 48hrs with DZA. Seventy-two hrs after cells plating or 48hrs post-treatment with DZA (100µM), cells were harvested according to the protocol for LC-Q-Exactive Focus HILIC mode of VIB Metabolomics Core and submitted to analysis (VIB Metabolomics Core, Leuven, Belgium). Proteins from the same cell lysates were isolated and quantified by BCA analysis that was used for data normalization. Data are expressed as arbitrary units and normalized with protein concentration, for each condition.

### Real time cell proliferation

Two thousand cells were seeded into 96-well plates in 200 µL complete medium 24hrs before DZA and DLP treatment. At the time of treatment, plates were inserted into the Incucyte S3 (Essen BioScience) and left during the duration of the experiment. Pictures were taken every 4hrs at 10X lens and analysis has been performed with Incucyte S3 software by confluence mask analysis.

### Establishment of SAHH-deficient PK4A Mes cells

In 6-well plates 4.10^5^ HEK293T cells were seeded 24hrs before being co-transfected with *Ahcy* Crispr/Cas9 knock-out plasmid or with a Crispr/Cas9 control plasmid (GenScript, *Ahcy* CRISPR Guide RNA (crRNA 5)) both coding for the Cas9 nuclease, a target-specific 20 nt guide RNA (gRNA) and a puromycin resistance gene, and with pMD2.G, psPAX2 plasmids (Addgene), using TransIT-LT1 (Mirus). Lentiviral supernatants were collected 24 to 48hrs post-transfection, and were filtered with 0.45µm.

Mes PK4A and KIC-derived LM (KIC-LM) cells (5.10^5^) were seeded on 6-well plates 24hrs before infection with lentiviruses. Lentiviruses were added into cells with 8µg/mL of Polybrene. At confluence (48 to 72hrs post-infection according to each cell lines), cells were detached with diluted accutase and further selected and maintained under puromycin at 1 µg/mL. SAHH-deficient (def) PC Mes PK4A cells represent a polyclonal population of cells. SAHH-def#1 Mes PK4A cells represent a clone isolated following limited dilutions of the polyclonal population of SAHH-def PC Mes PK4A cells. Two clones SAHH-def #A and #B KIC-LM cells were obtained from limited dilutions of a polyclonal SAHH-def KIC-LM cells.

### GSH/GSSG Ratio

Twenty-five thousand cells per well were plated into 96-well plate in 100µL of complete medium. 24hrs after, complete medium was removed, cells were rinsed with PBS and 100µL of Hank’s Balanced Salt Solution (HBSS) was added with or without DZA 100µM. Cells were lysed 48hrs later and analyzed as described in GSH/GSSG-Glo Assay.

### Real-time quantification of mitochondrial and cytosolic ROS and calcium production

In Rev. Figures 4F and rev. Supp 4D, 4 000 cells per well were plated in 96-well plates in 200µL complete medium for 24hrs. The day after, plated cells were treated with complete medium and with indicated DZA, GSH or L-Homocysteine. 48hrs later, cells were incubated for 30 min at 37°C in the dark with 2.5µM of green CellROX probe (ThermoFisher Scientific), or 10µM of Dihydrorhodamine 123 (ThermoFisher Scientific) or for 45 min at 37°C in the dark with 2.5µM of X-Rhod-1 AM or 5µM of Rhod-2 AM in free-Ca^2+^ media (*i.e.* S-MEM) supplemented with 2mM of glutamine. Cytosolic and mitochondrial ROS and calcium probes were removed and complete medium with or without indicated DZA, GSH, L-Homocysteine or menadione were added for 10hrs for mitochondrial ROS and mitochondrial/cytosolic calcium and 24hrs for cytosolic ROS. For calcium measurements, 1mM of EGTA was added to the complete medium. Pictures were taken at 10X lens into the Incucyte S3 (Essen BioScience) device, and ROS and calcium probe intensity, respectively determined at 58hrs and 72hrs post-treatment, has been normalized by cell confluence at this time point and is expressed relative to the highest value of ROS or calcium in each experimentation.

### Rhod-1-AM probe measurement

For Figure 4D and Supp 4C, 20 000 cells were seeded into 12-well plates in complete medium. DZA was added at indicated time. After maximum of 36hrs of drug stimulation, cells were removed with diluted accutase, rinsed with PBS and incubated with 2.5µM of X-Rhod-1-AM 30 min at 37°C in the dark in FACS buffer (PBS, BSA 5%, EDTA 5mM, no calcium). Cells were centrifuged, rinsed with PBS and resuspended in FACS buffer. Cells were analyzed by Flow cytometry (MACSQuant VYB flow cytometer, Miltenyi).

For immunofluorescent (IF) measurement, 20 000 cells were seeded in 12-well plates in complete medium directly on glass slides. DZA was added at indicated time. After maximum of 36hrs of drug stimulation, cells were incubated with 2.5µM of X-Rhod-1-AM 30 min at 37°C in the dark in FACS buffer. Cells were rinsed with PBS and fixed with 1% (wt/vol) formaldehyde for 30 min at room temperature. Cells were rinsed with PBS and slides were mounted with ProLong Dapi and analyzed by microscopy (Zeiss, Zeiss Axio Imager Z2+Apotome 2 SVO).

### RNA-Seq analysis

Mes PK4A cells (1.10^6^) were seeded into 100 mm dishes in 10 mL of complete medium. Cells were treated with DZA 100µM for 48hrs and then RNA extraction was performed according to Qiagen protocol. RNAs were quantified and RNA Integrity Numbers (RIN) were measured. RNAs with a RIN>9 were used. For RNA-Seq analysis (IntegraGen, Evry, France), the Bioconductor edgeR package was used to import raw counts into R statistical software. Differential expression analysis was performed using the Bioconductor limma package and the voom transformation. To improve the statistical power of the analysis, only genes expressed in at least one sample (FPKM ≥ 0.1) were considered. A p-value threshold of ≤ 0.05 was used to define differentially expressed genes between control and DZA treated samples. Gene list from the differential analysis was ordered by decreasing p-value.

### Cytoscape analysis

On the significant genes identified in the RNA-Seq analysis (p-value ≤ 0.05) gene set enrichment analysis was performed by clusterProfiler:GSEA function using the fgsea algorithm. Gene sets from MSigDB v7.2 database were selected among the C4_computational, C5_GO, C6_oncogenic, Hallmark classes, keeping only gene sets defined by 10-500 genes. Enrichment maps were constructed for gene sets that differed significantly between non-treated and DZA treated Mes PK4A cells with a FDR ≤ 0.10 and an overlap coefficient ≥ 0.25 and only related GO were represented. For Connectivity Map 2.0 analysis genes that were upregulated (red) and downregulated (blue) between NT vs DZA Mes PK4A cells were evaluated. Each GO term is represented by a dot and contain overexpressed (red dot) or downregulated (blue dot) genes in DZA treated cells compared to NT cells. Go terms are then merged into nodes with other associated terms.

### HeatMap

HeatMap of our experimentations have been designed using the Morpheus software (https://software.broadinstitute.org/morpheus).

For Figure 4B, representative significant genes (adjusted p value ≤0.05) involved in calcium regulation, ER stress and EMT pathways are represented in each triplicate of NT and DZA treated Mes PK4A cells and classified according to their adjusted p-value.

### Human gene expression database analysis and statistical analysis

We gathered clinicopathological and gene expression data of clinical pancreatic carcinoma samples from 17 publicly available mRNA expression datasets (Supplementary Table 6). Merged data was composed of 938 PDA and 55 liver metastasis samples. The informed patients’ consent to participate were obtained by the respective authors, while ethical considerations were reviewed by the corresponding institutional boards.

The data analysis required pre-analytic processing. First, we separately normalized each DNA microarray-based data set: the Affymetrix dataset was normalized by applying RMA (Robust Multichip Average) with the non-parametric quantile algorithm to the raw Affymetrix data (Bioconductor and associated-packages, R software), while the non-Affymetrix datasets were normalized with quantile normalization procedure. Then, Agilent or Affymetrix hybridization probes were mapped across different technological platforms (Stanford Online Universal Resource for Clones and ESTs (SOURCE, https://source-search.princeton.edu/), NCBI EntrezGene (, Homo sapiens gene information db, release from 04/27/2017) and Affymetrix NetAffx Annotation files (www.affymetrix.com; release from 01/12/2008)). When multiple probes mapped to the same gene, we retained the most variant probe in a particular dataset. For the RNA-Seq data, we used the available normalized data. We then log_2_-transformed the normalized genomics data. Subsequently, we corrected the 17 studies for batch effects using z-score normalization. Briefly, for each separate *AHCY*, ITGAV, and *CDH2* gene expression value in each study, we subtracted the mean of the gene in that dataset divided by its standard deviation, mean, and standard deviation being measured on primary cancer samples. All gene expressions in tumors were also measured as discrete values after comparison with median expression in the 993 samples: high expression was defined by value > median and low expression by value ≤ median. Genes from the methionine cycle and the transsulfuration pathway, (Supplementary Table 8) were similarly normalized and respective pathway metagenes were computed as the first component of a Principal Component Analysis (PCA) for each gene list. The Collisson’s molecular subtypes^51^ were applied as previously reported^52^.

A total of 755 PDA samples were informative for overall survival (OS). OS was calculated from the date of diagnosis to the date of death. Follow-up was measured from the date of diagnosis to the date of last news for event-free patients. Survivals, calculated using the Kaplan-Meier method, were compared with the log-rank test. All statistical tests were two-sided at the 5% level of significance. Statistical analysis was done using the survival package (version 3.1-12) in the R software (version 3.5.2; http://www.cran.r-project.org/).

### Human samples

Paraffine embedded samples of PDA (n=3) and corresponding liver metastasis (n=3), were obtained from patients with metastatic PDA who underwent surgery between 2009 and 2010. Histologic examination confirmed diagnosis of metastatic PDA. Tumor staging was performed according to the International Union Against Cancer TNM system. Before surgery, all patients had signed an informed consent form that had been approved by the local ethics committee (agreement reference of tissue collection: DC-2013-1857).

### Co-Immunofluorescence

Formalin-fixed paraffin-embedded, 5 µm-thick mouse PDA sections were deparaffinized in xylene and rehydrated through a series of graded ethanol concentrations. Antigen retrieval was performed in citrate buffer pH6/Tween 0.05%. PDA sections were rinsed 3 time with PBS and incubated 1hr at room temperature with 10% Serum/0.3% Triton. Tissue sections were rinsed 3 times with PBS and primary antibodies were incubated overnight, at 4°C in 1% BSA/0.05% Tween 20. Tissue sections were rinsed 3 times with PBS and labelled-secondary antibodies were incubated 1hr in the dark in 1% BSA/0.05% Tween 20. Tissue sections were rinsed 3 times with PBS and then with water. For slide mounting ProLong Dapi was used and slides were analyzed by microscopy (Zeiss, Zeiss Axio Imager Z2+Apotome 2 SVO).

### CFSE assay

PK4A cells (20 000) were counted and incubated with 5 µM of Cell Trace Violet in complete medium 20 min at 37°C in the dark. Cells were centrifuged, rinsed with PBS and plated in 12-well plate 24hrs before treatment with indicated concentration of DZA for 48hrs. Cells were removed with diluted accutase, rinsed with PBS and then resuspended in FACS buffer, and analyzed by flow cytometry (MACSQuant VYB flow cytometer, Miltenyi).

### Ahcy Activity Assay

Twenty-five thousand cells were plated into 96-well plate in 100µL of complete medium, 24hrs before DZA addition. After 48hrs treatment, cells were lysed and analyzed as described in Adenosylhomocysteinase (AHCY) activity fluorometric assay kit (Sigma-Aldrich).

### Statistical analysis

For *in vitro* experiments, we used Wilcoxon test for comparing 2 experimental groups in fold change or unpaired/paired two-tailed t-test for comparing 2 experimental groups. We used 1-way ANOVA followed by Sidak’s or Tukey correction for comparing more than 2 groups with 1 factor. For *in vitro* experiments, we used 2-way ANOVA followed by Bonferroni’s correction for comparing more than 2 experimental groups with 2 factors and 2-way repeated ANOVA followed by Bonferroni’s correction for time dependent experimentations. When, more than 2 groups are compared to a specific condition, we used a 2-way ANOVA followed by Dunnett’s correction. For *in vivo* experiments, to compare 2 experimental groups, we used Mann-Withney test. For more than 2 experimental groups, we used 2-way ANOVA followed by Bonferroni’s correction.

### Reporting summary

Further information on research design is available in the Nature Portfolio Reporting Summary linked to this article.

### Data availability

Publicly available databases used in this study are accessible through information mentioned in Table 6 with GEO database numbers and references. Source data are provided with this paper. All other data or unique materials used in this work are available from the corresponding authors on reasonable request.

### Reagents or Tools Table

| Reagents or Tools | Source | Identifier |
|---|---|---|
| **Antibodies** |  |  |
| Mouse anti-β-Actin monoclonal antibody (WB) | Sigma-Aldrich | Cat# A5316, RRID: AB_476743 |
| Mouse anti-β-Tubulin monoclonal antibody (WB) | Sigma-Aldrich | Cat# T8328, RRID: AB_1844090 |
| Rabbit anti-BIP monoclonal antibody (WB) | Cell Signaling Technology | Cat# 3177, RRID: AB_2119845 |
| Rabbit anti-CAMKK2 polyclonal antibody (WB) | ThermoFisher Scientific | Cat# 16910, RRID: AB_2798771 |
| APC Rat anti-CD24 monoclonal antibody (FACs) | BioLegend | Cat# 138505, RRID: AB_2565650 |
| APC/Cyanine 7 Rat anti-CD44 monoclonal antibody (FACs) | BioLegend | Cat# 103028, RRID: AB_830785 |
| Rabbit anti-CD51 monoclonal antibody (IHC-IF-WB) | Abcam | Cat# PA5-102029, RRID: AB_2851460 |
| PE Rat anti-CD51 monoclonal antibody (FACs) | BioLegend | Cat# 104105, RRID: AB_313074 |
| BV786 Rat anti-CD90 monoclonal antibody (FACs) | BD Biosciences | Cat# 740916, RRID: AB_2740558 |
| Pacific Blue Rat anti-CD106 monoclonal antibody (FACs) | BioLegend | Cat# 105722, RRID: AB_2304290 |
| Rabbit anti-CD106 monoclonal antibody (WB) | Abcam | Cat# ab134047, RRID: AB_2721053 |
| Goat anti-CK19 polyclonal antibody (WB) | Santa Cruz Biotechnology | Cat# sc-33111, RRID: AB_2234419 |
| Mouse anti-CTH monoclonal antibody (IHC) | Santa Cruz Biotechnology | Cat# sc-374249, RRID: AB_10986271 |
| Rabbit anti-CTH polyclonal antibody (WB) | Proteintech | Cat# 12217-1-AP, RRID: AB_2087497 |
| Mouse anti-DNMT1 monoclonal antibody (WB) | Novus | Cat# NB100-56519, RRID: AB_2093819 |
| Rat anti-E-Cadherin monoclonal antibody (WB) | Innovative Research | Cat# 13-1900, RRID: AB_86571 |
| Rabbit anti-EpCam polyclonal antibody (IHC) | Abcam | Cat# ab71916, RRID: AB_1603782 |
| Brillant Violet 605 Rat anti-EpCam monoclonal antibody (FACs) | BioLegend | Cat# 118227, RRID: AB_2563984 |
| Rabbit anti-GCLC polyclonal antibody (WB) | Proteintech | Cat# 12601-1-AP, RRID: AB_2278734 |
| Rabbit anti-HERPUD1 polyclonal antibody (WB) | ThermoFisher Scientific | Cat# PA5-29469, RRID: AB_2546945 |
| Rabbit anti-IRE1α monoclonal antibody (WB) | Cell Signaling Technology | Cat# 3294, RRID: AB_823545 |
| Rat anti-KI67 monoclonal antibody (IHC) | BioLegend | Cat# 652402, RRID: AB_11204254 |
| Rabbit anti-MAT1-A polyclonal antibody (WB) | Proteintech | Cat# 12395-1-AP, RRID: AB_2266186 |
| Rabbit anti-N-Cadherin polyclonal antibody (IHC-IF-WB) | Abcam | Cat# ab18203, RRID: AB_444317 |
| Mouse anti-SAHH monoclonal antibody (IHC-IF) | Santa Cruz Biotechnology | Cat# sc-271389, RRID: AB_10612213 |
| Rabbit anti-SAHH polyclonal antibody (WB) | Origene | Cat# TA332593 |
| Rabbit anti-SLUG monoclonal antibody (WB) | Cell Signaling Technology | Cat# 9585, RRID: AB_10330705 |
| Rabbit anti-SNAIL monoclonal antibody (WB) | Cell Signaling Technology | Cat# 3879, RRID: AB_2255011 |
| Rabbit anti-TRPM4 polyclonal antibody (WB) | Sigma-Aldrich | Cat# SAB2102584, RRID: AB_10606822 |
| Mouse anti-Vimentin monoclonal antibody (WB) | Santa Cruz Biotechnology | Cat# sc-32322, RRID: AB_628436 |
| **Chemicals, Peptides and Recombinant Proteins** |  |  |
| 3-Deazaadenosine (DZA) | Sigma-Aldrich | Cat# D8296 |
| GSH | Bertin Technologies | Cat# 14953 |
| L-Homocysteine | Santa Cruz Biotechnology | Cat# 280883 |
| Menadione | Sigma-Aldrich | Cat# M5625 |
| Mitomycin C | Sigma-Aldrich | Cat# 10107409001 |
| Polybrene | Sigma-Aldrich | Cat# TR-1002-G |
| Recombinant Human TGF-β1 | Peprotech | Cat# 100-21 |
| Sytox Green Ready Flow Reagent | ThermoFisher Scientific | Cat# R37168 |
| *Trans*IT-LT1 Transfection Reagent | Mirus | Cat# MIR 2300 |
| **Critical Commercial Assays** |  |  |
| Adenosylhomocysteinase (AHCY) activity fluorometric assay kit | Sigma-Aldrich | Cat# EPI021 |
| CellROX green reagent | ThermoFisher Scientific | Cat# C10444 |
| CellTraceTM CFSE cell proliferation kit | ThermoFisher Scientific | Cat# C34554 |
| Citrate buffer pH6 | Diapath | Cat# 70050 |
| GSH/GSSG-Glo Assay | Promega | Cat# V6611 |
| Liquid DAB and substrate chromogen system | DAKO | Cat# K3468 |
| RNeasy Micro Kit | Qiagen | Cat# 74134 |
| Target retrieval solution at pH 6 or 9 | DAKO | Cat# S1699 and #2367 |
| Tumor Dissociation Kit, mouse | Miltenyi Biotec | Cat# 130-096-730 |
| VECTASTAIN ABC kit | Vector | Cat# PK-6100 |
| Dihydrorhodamine 123 | ThermoFisher Scientific | Cat# D23806 |
| Rhod-2, AM | ThermoFisher Scientific | Cat# R1244 |
| X-Rhod-1-AM | ThermoFisher Scientific | Cat# X14210 |
| **Deposited Data** |  |  |
| Primary tumor/Liver metastasis RMA data: Expression data from 4 pairs of PDA and matched liver metastasis removed from 9-week-old KIC mice by laser capture microdissection | https://www.ncbi.nlm.nih.gov/geo/query/acc.cgi?acc=GSE266381 | GSE266381 |
| **Experimental Models: Cell Lines** |  |  |
| PK4A mouse cells | *Guillaumond et al (2013)* | RRID: CVCL_WB21 |
| **Experimental Models: Organisms/Strains** |  |  |
| Pdx1-Cre; Ink4a/Arf<sup>fl/fl;</sup>LSL-Kras<sup>G12D</sup>mice | Aguirre et al (2003) | https://doi.org/10.1101/gad.1158703 |
| **Recombinant DNA** |  |  |
| AHCY CRISPR Guide RNA 5 | GenScript |  |
| Control CRISPR Guide RNA 5 | GenScript |  |
| pMD2.G plasmid | Addgene | Plasmid# 12259 |
| psPAX2 plasmid | Addgene | Plasmid# 12260 |
| **Software and Algorithms** |  |  |
| Black Zen software | Zeiss | RRID: SCR_018163 |
| Cytoscape | Cytoscape | RRID: SCR_003032 |
| FlowJo Version 10 | FlowJo LLC | RRID: SCR_008520 |
| Image J | National Institutes of Health | RRID: SCR_003070 |
| Morpheus | Broad Institute | RRID: SCR_017386 |
| Prism Version 5.03 | Graph Pad | RRID: SCR_002798 |
| **Other** |  |  |
| Antibiotic/antimycotic solution | ThermoFisher Scientific | Cat# 15240062 |
| Bradfor Protein Assay | Bio-Rad | Cat# 5000205 |
| DMEM medium without glutamine (Gln-) | ThermoFisher Scientific | Cat# 11960-044 |
| Foetal bovine serum | BioSera | Cat# FB-1280 |
| gentleMACS C tubes | Miltenyi Biotec | Cat# 130-093-237 |
| Hank’s Balanced Salt Solution (HBSS) | ThermoFisher Scientific | Cat# 14025092 |
| L-glutamine (200 mM) | ThermoFisher Scientific | Cat# 25030024 |
| Protease inhibitor panel | Sigma-Aldrich | Cat# INHIB1 |
| Puromycin | ThermoFisher Scientific | Cat# A1113802 |
| StemPro Accutase cell dissociation reagent | ThermoFisher Scientific | Cat# A1110501 |
| Western chemiluminescent HRP substrate | Millipore | Cat# WBKLS0100 |

## ACKNOWLEDGEMENTS

We thank the PRIMACEN platform, University of Rouen (Rouen, France), with the collaboration of Magalie Bénard, the cell culture platform (PCC, CRCM, Marseille, France), the mice colony facility (PSEA, CRCM, Marseille, France), the bioinformatic platform (Cibi, CRCM, Marseille, France), the cytometry platform (CIPHE, Marseille, France) for technical assistance, and the VIB Metabolomics Core (Leuven, Belgium) for metabolomic analysis. E.J.P. was supported by the “Fondation pour la Recherche Médicale” (FRM). E.M. was supported by the French Ministry for Research and Education (MENRT) and “Ligue nationale contre le cancer” (LNCC). V.G. was supported by the LNCC and the “Association pour la Recherche sur le Cancer” (ARC). G.E. was supported by the FRM and S.D.B. was supported by “Fondation de France”. SV acknowledges supports from the National Institute of Cancer (INCa, Pair Pancreas 2018-082), the FRM, the ARC (PGA 2020), LNCC as “équipe labéllisée Ligue contre le cancer”, and the Cancéropôle PACA.

## AUTHORS CONTRIBUTIONS

E.J.P., E.M., V.G., G.E., S.D.B., performed the experiments. E.J.P. performed and assisted tracing experiments and GC/MS analysis. Q.D.C. analyzed the mouse transcriptomic data. E.L. contributed to the analysis of transcriptomic data. P.F., F.B., E.Ma., provided and analyzed human transcriptomic data sets. C.N., provided PDA patient samples. P.G., assisted cell sorting. N.D., provided primary human organoids. S.V., E.J.P., F.G., R.T., conceptualized the study. F.G., R.T., provided advice regarding experiments and writing of the manuscript. S.V. directed the study. S.V., E.J.P., F.G., G.E., P.F., Q.D.C., F.B. wrote the manuscript. Funding acquisition: S.V.

## COMPETING INTERESTS

All authors declare no competing interests

## ADDITIONAL INFORMATION

Correspondence and requests for materials should be addressed to Sophie Vasseur.

---

**Figure S1.**
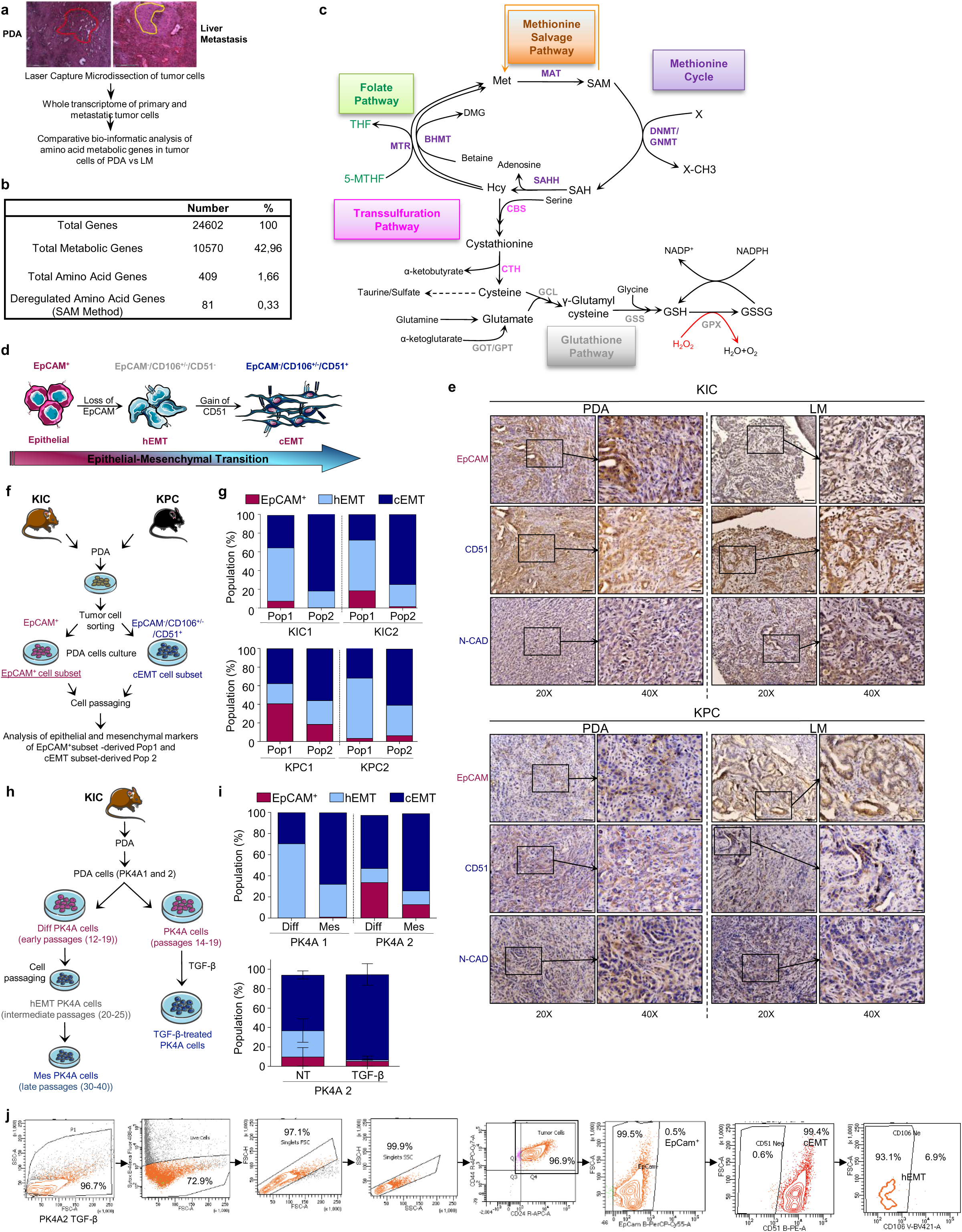
**a,** Schematic representation of the experimental approach to identify deregulated amino acid-associated metabolic genes in tumor cells from PDA (n=4 KIC mice) and liver metastasis (LM) (n=4 KIC mice), isolated with laser capture microdissection. **b,** Comparisons of LM (n=4) vs PDA (n=4) metabolic genes of KIC mice. 81 genes of the 409 amino acid genes are significantly upregulated. **c,** Schematic representation of the methionine cycle, the transsulfuration, and theglutathione pathways. Methionine (Met); Methionine Adenosyltransferase (MAT); S-Adenosylmethionine (SAM); DNA methyltransferase (DNMT); Glycine N-methyltransferase (GNMT); S-adenosyl-L-homocysteine (SAH); S-adenosylhomocysteine hydrolase (SAHH); Homocysteine (Hcy); Betaine-homocysteine S-methyltransferase (BHMT); N,N-dimethylglycine (Dmg); Methionine synthase (MTR); Tetrahydrofolate (THF); 5-methyltetrahydrofolate (5-MTHF); Cystathionine β-synthase (CBS); Cystathionine γ-lyase (CTH); Glutamate-cysteine ligase (GCL); Glutamic-oxaloacetic transaminase (GOT); Glutamate-pyruvate transaminase (GPT); Glutathione synthetase (GSS); Glutathione peroxidase (GPX); Glutathione (GSH); Glutathione disulfide (GSSG); Nicotinamide adenine dinucleotide phosphate (NADP/NADPH). **d,** Schematic representation of the expression of specific markers during EMT. The epithelial state is associated with EpCAM expression, while the hybrid state (hEMT) is characterized by loss of EpCAM and partial acquisition of CD106. The complete EMT state (cEMT) is associated with CD51 expression. **e,** Representative pictures of EpCAM, CD51 and N-cadherin immunostainings in PDA and LM of KIC (top, n=2 mice) and KPC (bottom, n=2 mice) mice (n=2 independent experiments). Scale bar: 20, 50 µM; 40X, 20 μm. **f,** Schematic representation of the experimental approach for KIC and KPC PDA cell isolation by fluorescence-activated cell sorting (FACS): epithelial (EpCAM^+^), and cEMT (EpCAM^-^/CD106^+/-^/CD51^+^) tumor cells were sorted, passaged, and further analyzed by flow cytometry and defined as Pop1 and Pop2, based on the expression levels of epithelial and mesenchymal markers. **g,** Flow cytometry analysis of epithelial and mesenchymal markers in Pop1 and Pop2 from KIC (n=2, top) and KPC (n=2, bottom) mice. Percentages of epithelial (EpCAM^+^), hybrid (hEMT) and complete (cEMT) EMT tumor cells subsets in Pop1 and Pop2 of each mouse are illustrated. **h,** Schematic representation of the experimental approach for the *in vitro* EMT induction in PK4A1 and 2 cells (2 independent batches). PK4A cells were isolated from the PDA of KIC mice and subjected to EMT by passaging of differentiated (Diff) PK4A cells from early passages (12-19), to intermediate passages (20-25, hEMT PK4A cells), until late passages (30-40, Mesenchymal (MesPK4A cells), or by treatment with TGF-β. **i,** Flow cytometry analysis of epithelial and mesenchymal markers in Diff and Mes (top; n=2 independent experiments) and TGF-b treated (bottom) PKA cells. For TGF-β experiments, data are expressed as mean±SEM from 3 independent experiments. Percentages of EpCAM^+^ (epithelial), hEMT and cEMT tumor cell subsets in Diff, Mes, non-treated (NT) and TGF-b treated PK4A cells are illustrated. **j,** Example of strategy gating to identify the different stages of EMT. To determine the percentage of epithelial, hEMT and cEMT cells in each population, viable cancer cells were selected by negative staining of Sytox green and then gated according to their expression of EpCam. EpCam^+^ cells were annotated as the EpCam^+^ subset. The EpCam^-^population was gated based on the expression of CD51. EpCam^-^/CD51^+^ cells, either CD106 positive or negative (CD106^+/-^), were annotated as cEMT cells. Finally, EpCam^-^/CD51^-^cells were annotated as hEMT cells.

**Figure S2.**
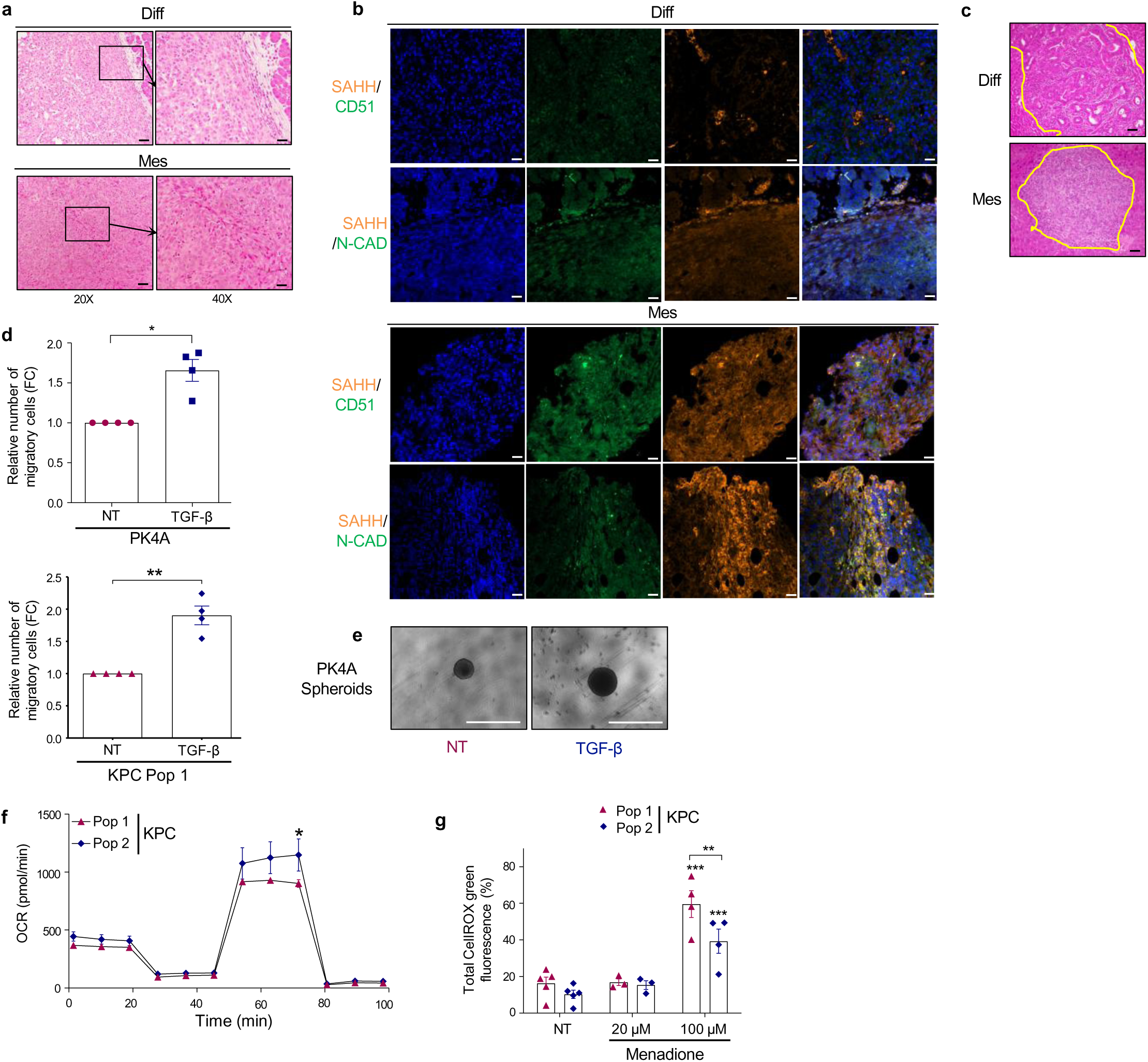
**a**, Representative pictures of HPS staining of Diff (n=6 mice, top) or Mes (n=6 mice, bottom) PDA. Scale bar: 20X, 50 µM; 40X, 20 μm. **b,** Representative images of SAHH (orange) /CD51 or N-cadherin (green) co-immunofluorescence on Diff (top) or Mes (bottom) PDA. Nuclei were visualized with DAPI (blue) (n=2 independent experiments). Co-localization was determined with z-stack analysis. Scale bar, 50 µM. **c,** Representative pictures of HPS staining of liver lobes of mice after Diff or Mes PK4A cell grafts in the spleen (n=5 mice/experimental group). Yellow lines delineate LM from liver tissue. Scale bar, 50 μm. **d,** Quantification of migration of NT and TGF-β-treated PK4A (top) and KPC Pop1 (bottom) cells after 12hrs. Relative values correspond to the FC in TGF-β-treated PK4A and KPC Pop1 cells relative to the respective NT cells (FC=1). Data expressed as mean±SEM from n=4 independent experiments. *\*p* < 0.05; *\*\*p* <0.01. Wilcoxon test. **e,** Representative images of NT and TGF-β-treated PK4A spheroids (n=3 independent experiments). Scale bar, 1 mm. **f,** Quantification of OCR in KPC Pop1 and Pop2 cells. OCR is expressed as mean±SEM from 3 independent experiments. Comparison Pop2 vs Pop1, *\*p* < 0.05; two-way ANOVA with Bonferroni’s correction. **g,** Total ROS production by control or menadione-treated KPC Pop1 and Pop2 cells shown as mean±SEM percentage of fluorescence from at least 3 independent experiments. Comparison Mena vs NT, *\*\*p* < 0.01; *\*\*\*p* <0.001; two-way ANOVA with Bonferroni’s correction.

**Figure S3.**
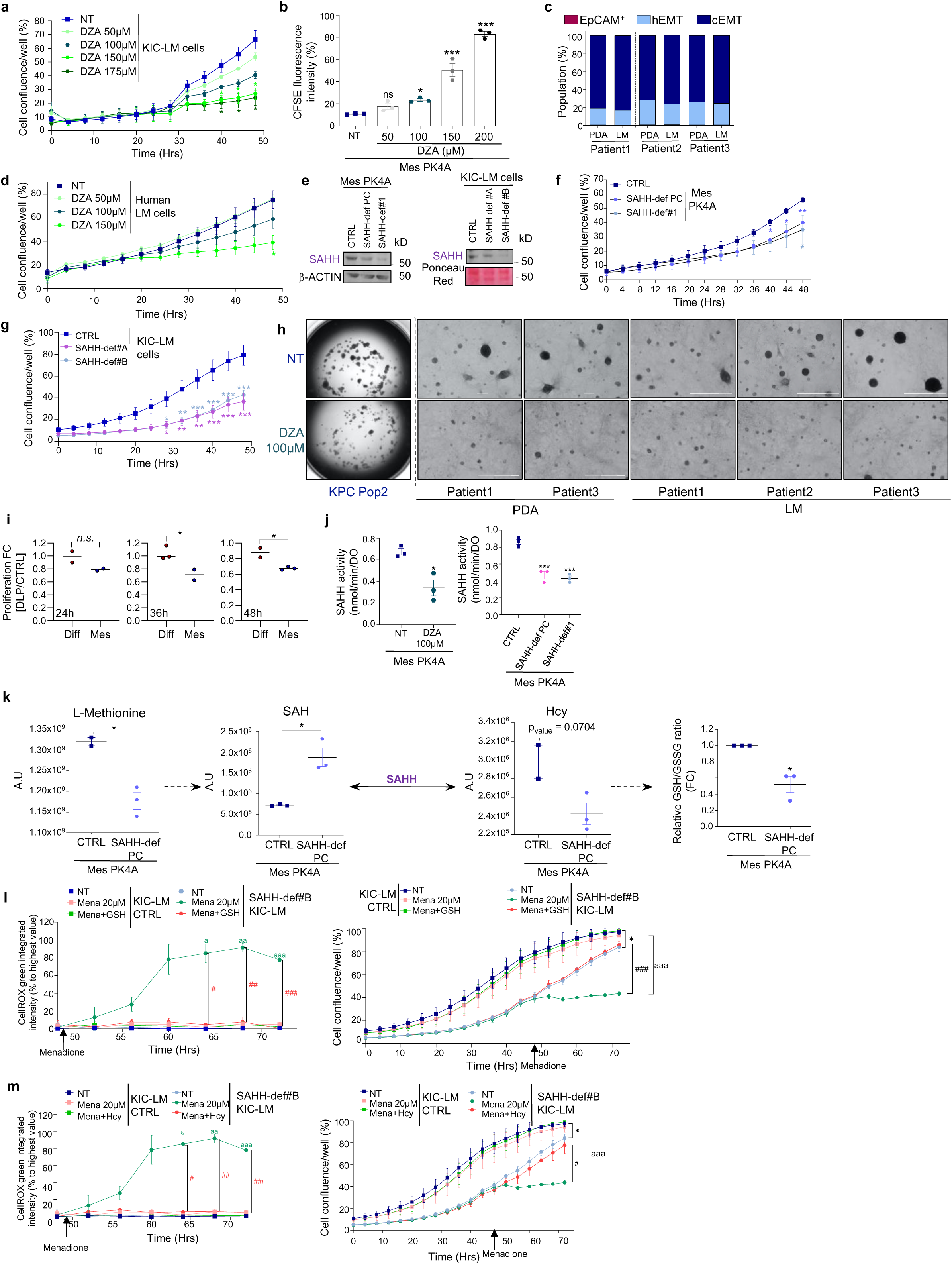
**a,** Proliferation of control or DZA-treated KIC-derived LM (KIC-LM) cellsover 50 hrs expressed as mean±SEM of cell confluence from 3 independent experiments. Comparison, DZA vs NT per time point. *\*p* < 0.05; repeated measures two-way ANOVA with Dunnett’s correction. **b,** Quantification of CFSE fluorescence in control or DZA-treated Mes PK4A cells represented as mean±SEM of the cell percentage from 3 independent experiments. Comparison DZA vs NT, *\*p* < 0.05, *\*\*\*p* <0.001, one-way ANOVA with Sidak’s correction. **c,** Flow cytometry analysis of epithelial and mesenchymal markers in human PDA-(n=3 patients: 1, 2, 3) or LM-(n=3 patients: 1, 2, 3) derived tumor cells, shown as the percentage of EpCAM^+^, hEMT and cEMT tumor cell subsets in each PDA or LM from patient 1, 2, and 3. **d,** Proliferation of control or DZA-treated human LM-derived tumor cells over 50 hrs expressed as mean±SEM of cell confluence from 4 independent experiments on n=3 human LM-derived tumor cells (patients 1, 2, and 3). Comparison, DZA vs NT per time point, *\*p* < 0.05, repeated measures two-way ANOVA with Dunnett’s correction. **e,** Levels of SAHH in CTRL, SAHH-def PC and SAHH-def#1 Mes PK4A cells (left) and in SAHH-deficient KIC-LM cells (SAHH-def #A and #B clones of KIC-LM cells, right). β-ACTIN (left) and ponceau red staining (right) were used as loading controls. Representative immunoblots from 3 independent experiments. **f,** Proliferation of CTRL, SAHH-def PC and SAHH-def#1 Mes PK4A cells expressed as mean±SEM of cell confluence from 3 independent experiments. Comparison, SAHH-def PC vs CTRL and SAHH-def#1 vs CTRL; *\*p* < 0.05, *\*\*p* < 0.01; repeated measures two-way ANOVA with Bonferroni’s correction. **g,** Proliferation of CTRL, SAHH-def#A and SAHH-def#B KIC-LM cells expressed as mean±SEM of cell confluence from 3 independent experiments. Comparisons, SAHH-def#A or SAHH-def#B vs CTRL per time point; *\*p* < 0.05, *\*\*p* < 0.01, *\*\*\*p* < 0.001; repeated measures two-way ANOVA with Bonferroni’s correction. **h,** Representative images of control or DZA-treated KPC Pop2 (left), human (right) PDA-(Patient 1, 3) and LM-(patient 1, 2, 3) organoids over 48h (n=3 experiments). Scale bar, 1 mm. **i,** Proliferation of Diff and Mes PK4A cells treated with 10 mM of DLP for 24, 36 and 48 hrs. Mean±SEM of the Fold change of the cell confluence relative to the non-treated cells at each time point and normalized to values at 0 hours from at least two independent experiments is shown. Comparison, Mes vs Diff PKA cells per time point; *\*p* < 0.05; two-tailed unpaired t test. **j,** Quantification of SAHH activity in control or DZA-treated Mes PK4A cells (left), and in CTRL, SAHH-def PC and SAHH-def#1 Mes PK4A cells (right) expressed as mean±SEM from3 independent experiments. *\*p* < 0.05, *\*\*\*p* <0.001; unpaired two-tailed t-test (right); one-way ANOVA with Sidak’s correction (left). **k,** Intracellular L-Methionine, SAH, and Hcy in CTRL and SAHH-def PC Mes PK4A cells (left). Data expressed in A.U. as mean±SEM of at least 3 biological replicates. *\*p* < 0.05; paired two-tailed t-test. The GSH/GSSG ratio measured in CTRL and SAHH-def PC Mes PK4A cells (right) is expressed as mean ±SEM FC relative to CTRL from 3 independent experiments. *\*p* < 0.05; Wilcoxon test. **l,** Total ROS production (left) and proliferation (right) for CTRL and SAHH-def#B KIC-LM cells treated with menadione (Mena) combined with 1 mM of GSH for 72hrs. ROS levels are expressed as the percentage of the highest ROS value and presented as mean±SEM from 3 independent experiments, while proliferation is expressed as mean±SEM of the cell confluence from 3 independent experiments. *Comparison between “CTRL NT” and “SAHH-def#B NT”; ^a^comparison between “SAHH-def#B+Mena” and “CTRL+Mena” cells; ^#^comparison between “SAHH-def#B+Mena” and “SAHH-def#B+Mena+GSH” cells. ^*,*#,a*^ *p* < 0.05; *^##,aa^p* <0.01; *^###,aaa^p* < 0.001; repeated measures two-way ANOVA with Dunnett’s correction (ROS levels); multiple unpaired t-tests at endpoint (proliferation). **m,** Total ROS production (left) and proliferation (right) for CTRL and SAHH-def#B KIC-LM cells treated with mena combined with 1 mM of Hcy for 72hrs. ROS levels are expressed as the percentage of the highest ROS value and presented as mean±SEM from 3 independent experiments, while proliferation is expressed as mean±SEM of the cell confluence from 3 independent experiments. *Comparison between “CTRL NT” and “SAHH-def#B NT”; ^a^comparison between “SAHH-def#B+Mena” and “CTRL+Mena” cells; ^#^comparison between “SAHH-def#B+Mena” and “SAHH-def#B+Mena+Hcy”. *^,*#,a*^*p* < 0.05; *^##,aa^,p* <0.01; *^###,aaa^p* < 0.001; repeated measures two-way ANOVA with Dunnett’s correction (ROS levels); multiple unpaired t-tests at endpoint (proliferation).

**Figure S4.**
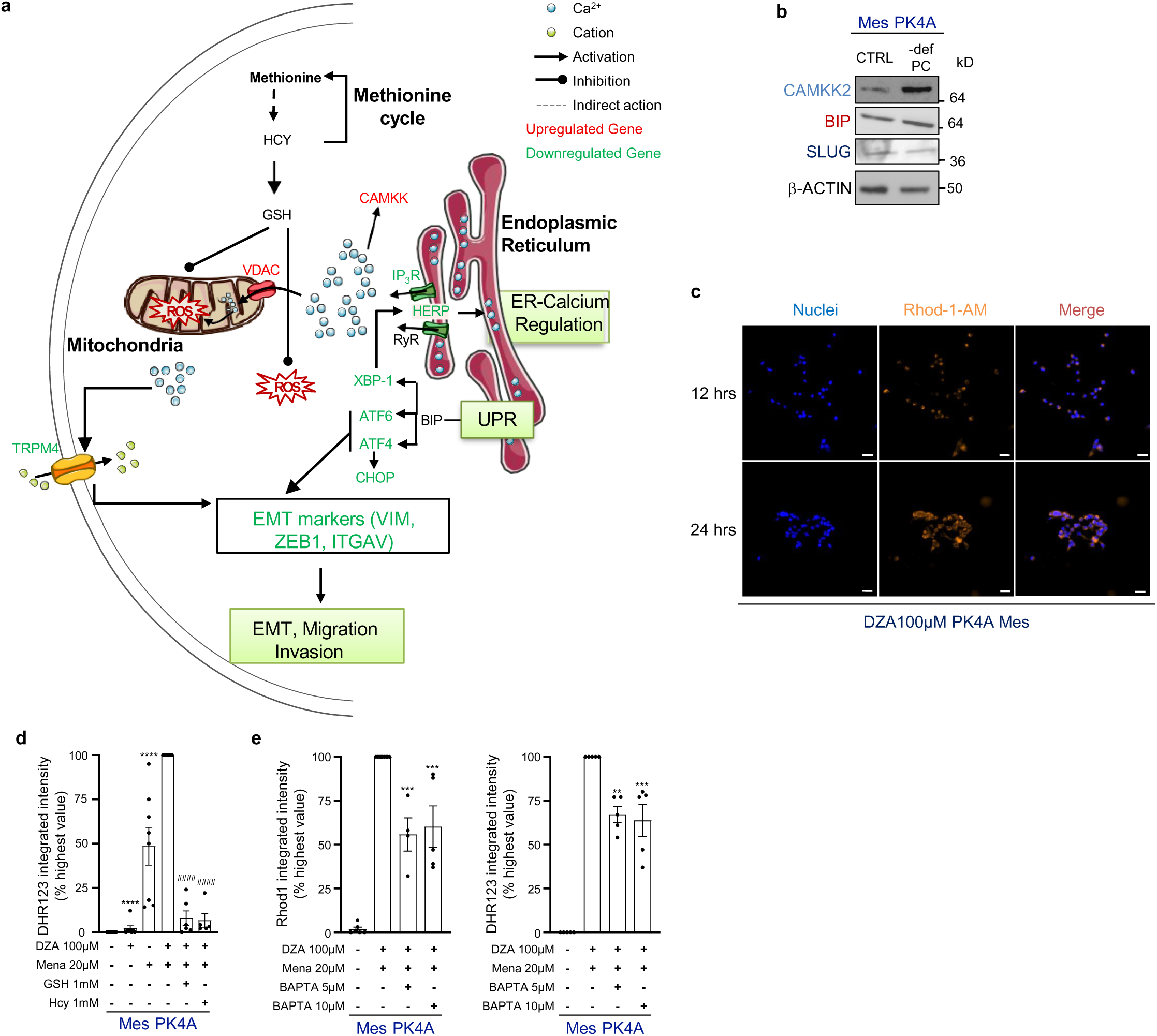
**a,** Schematic representation of the regulators of calcium homeostasis in the endoplasmic reticulum, the UPR (Unfolded Protein response), EMT, calcium flux in the cytosol and mitochondria, of the methionine cycle, and their cross-regulation. Genes are shown in red (upregulated) or green (downregulated) in DZA-treated compared to control tumor cells identified by RNAseq. **b,** Representative immunoblot of CAMKK2, BIP, and SLUG in CTRL and -def PC Mes PK4A cells from 3 independent experiments). β-Actin was used as loading control. **c**, Representative immunofluorescence images of Rhod-1-AM (orange) in control or DZA-treated Mes PK4A cells over 12 and 24hrs. Nuclei were visualized with DAPI (blue). n=3 independent experiments. Scale bar, 50µM. **d**, Quantification of DHR123 in Mes PK4A cells control or treated with DZA and Mena alone or combined, and with DZA+Mena combined with GSH or Hcy. Data expressed as mean±SEM percentage of the highest intensity value from 5-8 independent experiments. *\**Comparison between “Mena + DZA” *vs* “DZA” or “Mena”; ^#^comparison between “Mena+DZA+GSH or Hcy” and “Mena+DZA”. ****^,####^ *p* <0.0001. one-way ANOVA with Tukey’s correction **e.** Quantification of X-Rhod1-AM (left) and of DHR123 (right) in Mes PK4A cells control or treated with DZA. Menadione and BAPTA were added in the presence of EGTA 10 hrs before endpoint. Data are expressed as the percentage of the highest intensity value and presented as mean±SEM from 4-9 independent experiments. *Comparison between “Mena+DZA+BAPTA” vs “Mena+DZA”. **p <0.01, ***p <0.001; one-way ANOVA with Tukey’s correction.

**Figure S5.**
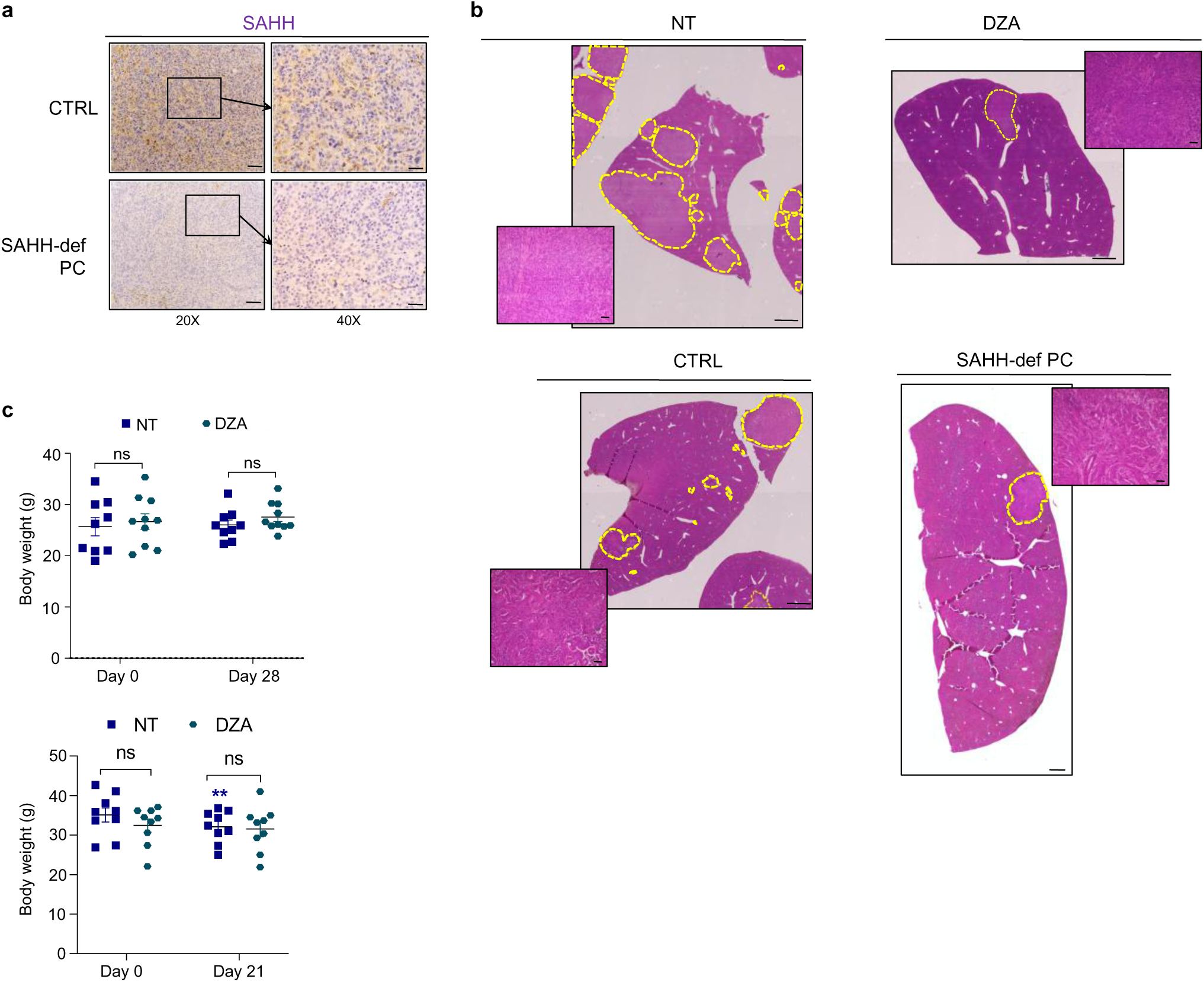
**a**, Representative images of SAHH immunostaining of CTRL or SAHH-def PC Mes PK4A tumors of KI mice 28 days post-implantation (n=3 independent experiments). Scale bars: 20X, 50 µM; 40X, 20 μm. **b,** Representative images of HPS staining of liver lobes of KI mice treated with vehicle (NT) (n=9 mice) or with DZA (n=10 mice) after implantation in the spleen of Mes PK4A cells (top), or of KI mice 28 days post-implantation in the spleen of CTRL or SAHH-def PC Mes PK4A cells (bottom) (n=5 mice each). Metastatic areas are distinguished from the liver parenchyma with yellow dotted lines. Scale bar, 1 mm; 50 μm for the insets. **c,** Body weight of KI mice orthotopically (top) or intrasplenically implanted (bottom) with Mes PK4A cells and treated with vehicle (NT) (n=9 mice) or DZA (n=10 mice) at day 0, 28 or 21 (end-point days). Data expressed as mean±SEM. *\*\*p* <0.01; two-way ANOVA with Bonferroni’s correction. *Comparison between NT Day 0 and NT Day 21.

